# Activated I-BAR IRSp53 clustering controls the formation of VASP-actin-based membrane protrusions

**DOI:** 10.1101/2022.03.04.483020

**Authors:** Feng-Ching Tsai, J. Michael Henderson, Zack Jarin, Elena Kremneva, Yosuke Senju, Julien Pernier, Oleg Mikhajlov, John Manzi, Konstantin Kogan, Christophe Le Clainche, Gregory A. Voth, Pekka Lappalainen, Patricia Bassereau

## Abstract

Filopodia are actin-rich membrane protrusions essential for cell morphogenesis, motility, and cancer invasion. How cells control filopodia initiation on the plasma membrane remains elusive. We performed experiments in cellulo, in vitro and in silico to unravel the mechanism of filopodia initiation driven by the membrane curvature sensor IRSp53. We showed that full-length IRSp53 self-assembles into clusters on membranes depending on PIP2. Using well-controlled in vitro reconstitution systems, we demonstrated that IRSp53 clusters recruit the actin polymerase VASP to assemble actin filaments locally on membranes, leading to the generation of actin-filled membrane protrusions reminiscent of filopodia. By pulling membrane nanotubes from live cells, we observed that IRSp53 can only be enriched and trigger actin assembly in nanotubes at highly dynamic membrane regions. Our work supports a regulation mechanism of IRSp53 in its attributes of curvature sensation and partner recruitment to ensure a precise spatial-temporal control of filopodia initiation.

## Introduction

Plasma membrane shaping relies on a precisely controlled coupling of the plasma membrane and the actin cytoskeleton. (*1–3*). A prominent example is filopodia, thin, finger-like membrane protrusions (typical diameter of 100-300 nm and length on the order of 10 μm) filled with parallel actin filaments bundled by fascin (typically 10–30 filaments) (*4–7*). Cells use filopodia to explore and sense their environment. Thus, filopodia are critical in numerous cellular processes, including polarized cell migration and adhesion, and in the tissue environment, embryogenesis, cancer invasion and cell-cell communication (*2, 6, 7*). Filopodia formation employs different mechanisms involving different sets of actin regulatory proteins (*6–8*). So far, two distinct mechanisms of filopodia generation have been proposed: the convergent elongation model relying on the reorganization of the pre-existing Arp2/3 complex-mediated branched actin network in lamellipodia (*9–11*), and the tip nucleation model in which de novo actin assembly is triggered by formin-family actin nucleators (*6*). The two mechanisms are not necessarily mutually exclusive and are most likely cell type dependent (*6, 12–14*). Notably, recent cell biology studies proposed another filopodia generation mechanism in which the membrane curvature sensing protein IRSp53 (insulin receptor substrate p53) forms local clusters on the plasma membrane to recruit the actin polymerase VASP (vasodilator-stimulated phosphoprotein), which promotes actin filament elongation at the onset of filopodia initiation (*15–18*). Yet, it remains poorly understood how cells control precisely when and where to trigger actin assembly on the plasma membrane to initiate filopodia (*6–8*).

IRSp53 (also known as BAIAP2), an inverse BAR (I-BAR, Bin-Amphiphysin-Rvs161/167) domain protein, is a crucial player in coordinating actin assembly and membrane dynamics in filopodia formation (*15–24*). IRSp53 features an N-terminal I-BAR domain (also known as IMD, IRSp53-MIM homology Domain), followed by a CRIB-PR (Cdc42/Rac interactive binding-proline rich) domain, and a canonical SH3 (Src homology 3) domain. Purified I-BAR domains spontaneously assemble into crescent-shaped dimers (*24*) that can bind to negatively charged lipids such as PS and PI(4, 5)P2 (hereafter as PIP2) (*25, 26*). The SH3 domain of IRSp53 allows it to interact with many actin regulators, such as VASP (*15, 16, 18*), Eps8 (*20*), N-WASP (*16*), WAVE2 (*27, 28*), and mDia1 (*28*). Importantly, it was shown that IRSp53 exhibits a closed, autoinhibited conformation due to the binding of its SH3 domain to its CRIB-PR motif (*29*). The most well-known activator of IRSp53 is Cdc42 (*15, 16, 19, 29*). Additionally, PIP2 and cytoskeleton effectors, such as Eps8 and VASP, have been shown to synergize with Cdc42 in IRSp53 activation to its fully open conformation (*19, 21, 29-31*). Importantly, it has been shown that 14-3-3 binds to phosphorylated IRSp53, keeping it in the inhibited state (*21, 30, 31*).

Biophysical studies in cellulo and in reconstituted systems have revealed that the I-BAR domain of IRSp53 can sense and generate similar negative membrane curvature that is found in filopodia (*26, 32–34*). Consistently, when overexpressing either the I-BAR domain or the full-length IRSp53 protein in cells, the generation of membrane protrusions reminiscent of filopodia were observed (*15-17, 19-23*). The majority of these I-BAR-induced protrusions contain actin filaments, albeit many of them have a low amount of actin (*17*). Notably, it was shown that full-length IRSp53 forms clusters on the plasma membrane that are followed by the accumulation of VASP for filopodial initiation (*15*). This tendency to cluster was supported by an in vitro study where, by pulling membrane tubes from giant unilamellar vesicles (GUVs) encapsulating the I-BAR domain, it was shown that at low concentrations and low curvature, the I-BAR domain phase separates along the tubes (*33*). Moreover, theoretical modeling has predicted how IRSp53 and actin cooperatively drive the formation of membrane protrusions (*35, 36*). Yet, little is known about how IRSp53 by itself clusters on the plasma membrane, and recruits VASP and actin. Moreover, we still lack a comprehensive mechanistic description of how IRSp53 and VASP cooperate with actin and its regulatory proteins to initiate filopodia.

VASP is a member of the Ena/VASP protein family that has been indicated to contribute to filopodia formation in cells (*6-9, 14, 37, 38*). VASP forms homo-tetramers that are weakly processive as actin polymerases. When forming oligomers or clusters, or when the actin filaments are bundled by fascin, VASP becomes more processive on the barbed ends of the actin filaments (*35, 39, 40*). Consistent with its actin polymerase activity, VASP clusters are part of the filopodial “tip complex”, where actin assembly occurs on the plasma membrane to elongate the filopodium (*9, 11, 41*). Both IRSp53 and another actin-regulator, lamellipodin, were shown to promote VASP clustering on the plasma membrane (*15, 18, 19, 22, 42, 43*). However, how VASP clusters facilitate filopodial initiation remains to be explored (*6, 7*).

To uncover how IRSp53, VASP and actin cooperate to initiate filopodia, we performed experiments in cellulo, in vitro, and in silico. Using in vitro reconstitution systems, we demonstrated that on PIP2-membranes, purified full-length IRSp53 self-assembles into clusters that are crucial for the recruitment of VASP to locally trigger actin assembly, giving rise to filopodia-like membrane protrusions. Our coarse-grained simulations showed that PIP2 is key for IRSp53 clustering. Our in vitro assays revealed that fascin is not required for filopodia initiation; however, we observed that in live cells, fascin enhances filopodia elongation and stability. Finally, to unravel the regulation of IRSp53 activity in live cells, we performed an in cellulo biophysical assay in which membrane nanotubes having a membrane geometry reminiscent of cellular filopodia, but initially devoid of actin, are pulled from the cells using optical tweezers. This assay allows us to reveal the regulated activation of IRSp53 in cells by detecting two of its functions: its ability to sense the negative membrane curvature of the pulled nanotubes and to trigger actin assembly inside the nanotubes. We found that IRSp53 is active only in membrane nanotubes that are pulled from regions of cell surfaces where the plasma membrane exhibits dynamic shape changes. Taken together, our results provide fundamental insights into how the curvature sensor protein IRSp53 synergizes with the actin polymerase VASP in filopodia initiation. Our findings further suggest that IRSp53 is tightly regulated in cells to have a rigorous control over filopodia initiation.

## Results

### IRSp53 self-assembles into clusters that recruit VASP on PIP2-membranes

Earlier work on MEF cells derived from IRSp53 null-mice provided evidence that IRSp53 is responsible for the formation of VASP clusters at the leading edge of migrating cells (*15*). It was shown that the accumulation of IRSp53 precedes the recruitment of VASP prior to the formation of filopodia (*15*). To further examine the roles of IRSp53 and VASP in filopodia initiation, we expressed IRSp53-eGFP and VASP-RFP in live Rat2 cells, which assemble frequent endogenous filopodia (*44*). By tracking their fluorescence signals at the plasma membrane, we found that IRSp53 is present along filopodia (Fig. 1A, brackets), while VASP tends to enrich at the filopodial tips (Fig. 1A, arrows). Importantly, before the appearance of filopodia, IRSp53 is enriched at the plasma membrane often as small clusters, while VASP displays quite a diffuse cytoplasmic localization with enrichment at focal adhesions (Fig. 1A). At the onset of a filopodium, the emergence of larger IRSp53 clusters occurs, and this is followed by the birth of VASP clusters (Fig. 1B and Movie S1). To reveal the evolution of IRSp53 and VASP clusters at the plasma membrane before and after filopodia formation, we generated adaptive kymographs with a previously developed automated method (*42*). The adaptive kymographs follow the movement of the plasma membrane (Fig. 1C) and map the signals of IRSp53 and VASP into vertical lines to create space-time plots (Fig. 1D). The kymographs confirm that IRSp53 clustering at the plasma membrane precedes VASP accumulation and filopodia initiation (Fig. 1D). These data, together with the earlier work (*15–18*), provide evidence supporting a previously proposed mechanism in which IRSp53 clustering at the plasma membrane induces VASP recruitment to locally polymerize actin on the membrane for driving protrusion formation (*15–18*). In this scenario, the synergistic functions of IRSp53, VASP, actin, and the filopodia-specific actin filament bundling protein, fascin, would be sufficient to drive protrusion formation.

**Fig 1.**
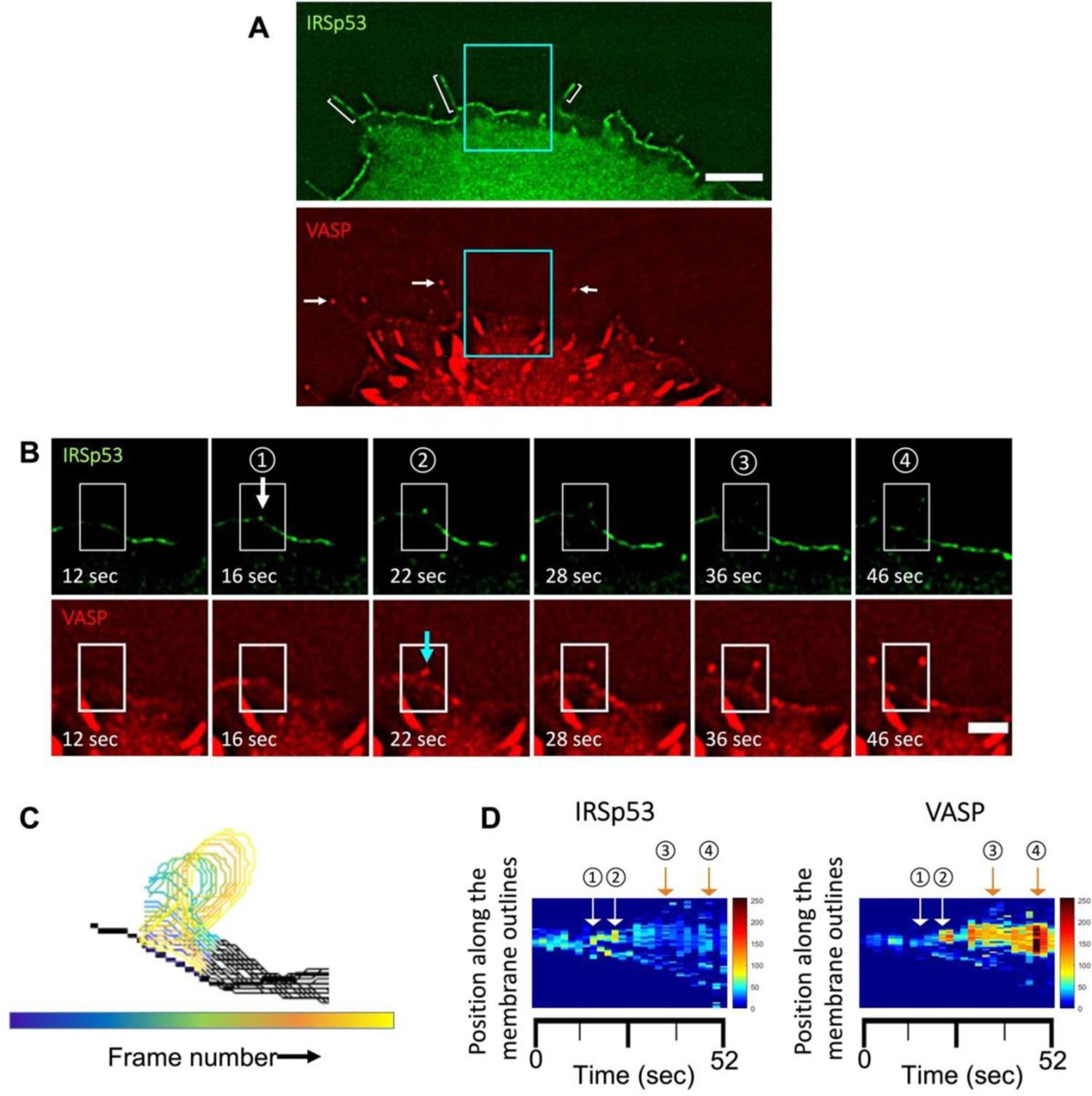
Dynamics of VASP clusters assembled from preexisting IRSp53 clusters on the plasma membrane in filopodia initiation. (A) Widefield fluorescence image of a representative Rat2 cell transfected with pEGFP-N1-IRSp53L (IRSp53-eGFP) and pmTagRFP-VASP (VASP-RFP). Brackets indicate some filopodia where IRSp53 is present along them. White arrows indicate the same filopodia to demonstrate that VASP is enriched in their tips. Scale bar, 5 μm. (B) Time-lapse images of a filopodium formation. Magnification of the indicated area shown in (A), cyan boxes. The white arrow indicates the appearance of a IRSp53 cluster followed by a VASP cluster indicated by a cyan arrow in the onset of filopodia formation. White boxes indicate the selected area used to generate outlines of plasma membrane positions over time shown in (C). Scale bar, 2 μm. (C) Colored outlines of membrane positions in the region indicated by the white boxes shown in (B). Total 27 frames, frame interval 2 sec. (D) Adaptive kymograph maps replot the detected membrane profiles in (C) in the y-axis and the corresponding time points in the x-axis to show the dynamics of IRSp53 (*Left*) and VASP (*Right*) on the plasma membrane over time. Y-axis shows the membrane positions of the proteins, and the x-axis shows the time (in second, total 27 frames). Color maps: low fluorescence intensity in blue, and high fluorescence intensity in red. Circled numbers correspond to the frames indicated in (B).

To test this hypothesis in a better-controlled experimental environment than in live cells, we developed an in vitro reconstitution system composed of giant unilamellar vesicles (GUVs) as model membranes, and purified IRSp53, VASP, actin and fascin. We first assessed if IRSp53 can self-assemble into clusters on model membranes without the presence of other proteins. We purified full-length human IRSp53 and labelled it with AX488 dyes (AX488-IRSp53) for visualization by fluorescent confocal microscopy. We generated GUVs containing 5 mole% of PIP2, given that it is the key phosphatidylinositol for an IRSp53-membrane interaction and thus necessary for filopodia generation (*26*). We used IRSp53 at 16 nM in its dimer form (i.e., 32 nM as a monomer) to be comparable to the cellular concentration of IRSp53 (29.7 nM to 453 nM, in which 29.7 nM is sufficient for filopodia generation)(*16*). Interestingly, by incubating AX488-IRSp53 with PIP2-GUVs, we observed clusters of IRSp53 on the GUV membranes (72%, N = 58 GUVs (technique replicates), n = 3 independent sample preparations (biological replicates) (*45*)) (Fig. 2A, arrows). These clusters are reminiscent of IRSp53 clusters observed in cells at the onset of filopodia formation. Additionally, we observed that a small fraction of GUVs (usually less than 10% of the whole GUV population) have inward membrane protrusions, where IRSp53 is in the interior of the protrusions (Fig. 2A arrowhead, Fig. S1, and Movie S2). This observation demonstrates that the full-length IRSp53 is functional such that it can generate negative membrane curvature.

**Fig. 2.**
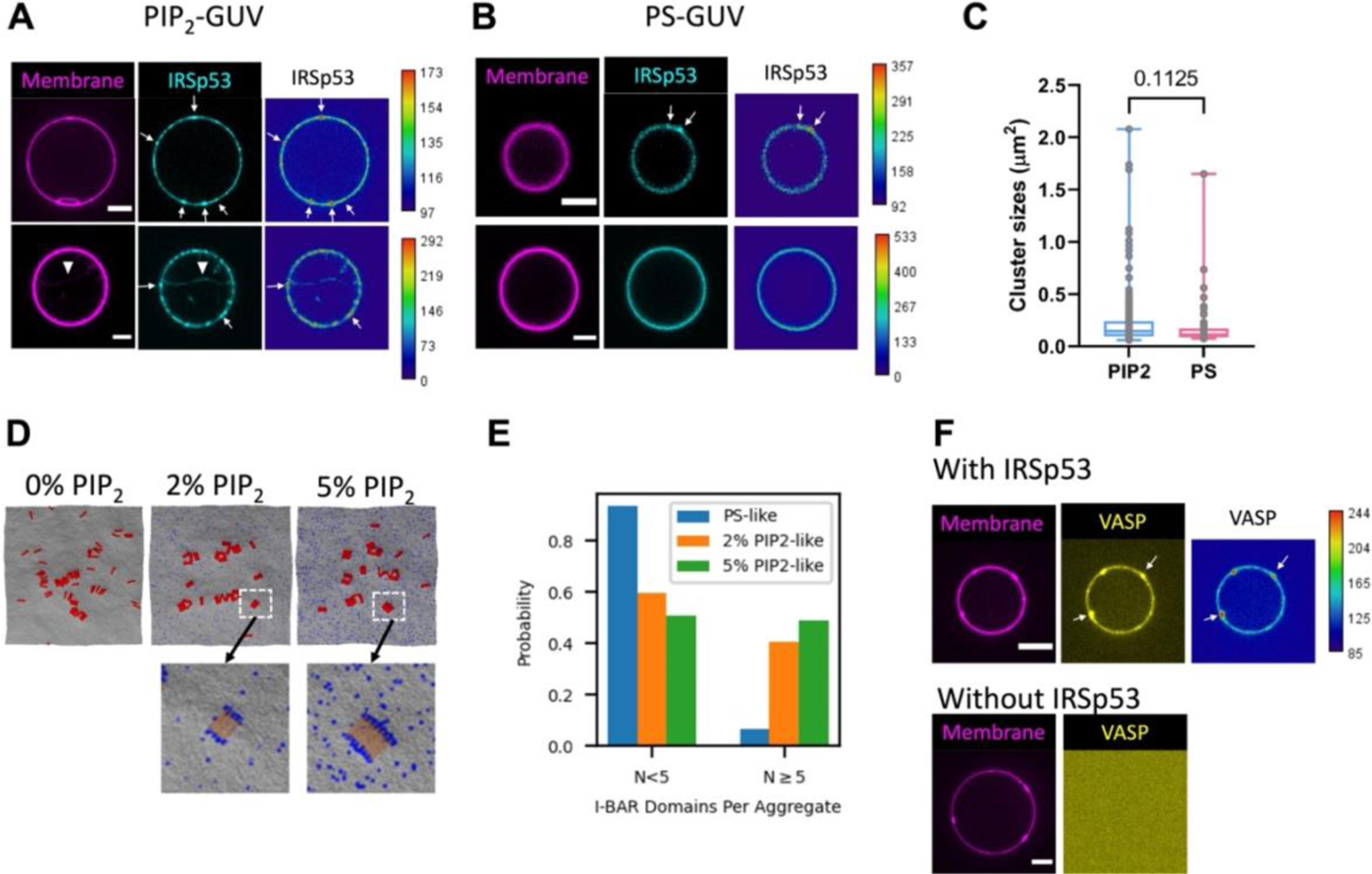
IRSp53 self-assembles into clusters and recruits VASP on PIP_2_-membranes. (A) and (B) Representative confocal images of GUVs incubated with 16 nM of AX488 labeled full-length IRSp53. In (A) GUVs contain total brain lipid extract (TBX) supplemented with 0.5% BODIPY TR ceramide (TR-ceramide) and 5% PIP_2_. In (B) GUVs contain TBX supplemented with 0.5% TR-ceramide and 25% dioleoyl-phosphatidylserine (DOPS). Color codes: magenta, TR-ceramide and cyan, IRSp53. White arrows indicate some of the IRSp53 clusters on the GUV membranes. Arrowhead indicates an inward membrane tube generated by IRSp53. Heat maps: low IRSp53 fluorescence intensity in blue, and high IRSp53 fluorescence intensity in red. (C) Sizes of IRSp53 clusters on PIP_2_-GUVs and on PS-GUVs. Each gray data points represent one cluster. PIP_2_-GUV: total 225 clusters, N = 42 GUVs, n = 3 sample preparations. PS-GUV: total 55 clusters, N = 12 GUVs, n = 2 sample preparations. Statistical test: two-tailed Mann-Whitney test, *p* = 0.1125. (D) (*Top*) Representative snapshots of coarse-grained (CG) simulations from PS-like (i.e., 0% PIP_2_-like), 2% PIP_2_-like, and 5% PIP_2_-like membranes. Membrane CG beads are shown in grey, PIP_2_-like CG beads in blue, and I-BAR domains in red. (*Bottom*) Enlarged areas as indicated by the white boxes in 2% and 5% PIP_2_-like membranes. Note that the color of the I-BAR domain is in light yellow to enhance the visibility of the PIP_2_ clusters, shown in blue, at the peripheries of the I-BAR domain clusters. (E) Probability of I-BAR domain aggregate size to be < 5 molecules, or ≥5 for PS-like (0% PIP_2_), 2% PIP_2_-like and 5% PIP_2_-like membranes as shown in (D). (F) Representative confocal images of GUVs incubated with AX488 labelled VASP (shown in yellow) together with (*Top*) or not (*Bottom*) IRSp53 (unlabeled). Protein concentrations: IRSp53, 16nM (dimer) and VASP, 4 nM (tetramer). GUVs contain TBX supplemented with 0.5% TR-ceramide (shown in magenta) and 5% PIP_2_. Color maps in (A) for IRSp53 signals, (B) for ISRp53 signals and (F) for VASP signals: low fluorescence intensity in blue, and high fluorescence intensity in red. Scale bars, 5 μm.

To explore the role of PIP2 in IRSp53 clustering, we tested if IRSp53 could form clusters upon interacting with another negatively charged lipid, PS, given that IRSp53-PIP2 binding is driven by electrostatic interactions (*26*). We replaced PIP2 with PS (25 mole%, PS has a net charge of −1) on GUVs while keeping the amount of negative charges comparable to that of PIP2-GUVs (5 mole% of PIP2. At pH 7, the charge of PIP2 is expected to be around −4 (*46*)). We found that IRSp53 can form clusters also on PS-GUVs (Fig. 2B). The number of clusters per GUV are comparable on PIP2-GUVs and PS-GUVs (Fig. S2). However, the clusters are larger on PIP2-GUVs as compared to PS-GUVs (Fig. 2C). To elucidate the mechanism underlying IRSp53-clustering on PIP2-membranes, we performed coarse-grained (CG) simulations. We generated PS-like and PIP2-like membrane sheets. The PS-like membrane is a quasi-monolayer of membrane beads that uniformly interacts with the I-BAR domain membrane binding surface. The PIP2-like membranes are nearly identical except a subset of membrane beads (2% or 5%) preferably interact with the ends of the I-BAR domains, as previously reported (*26*). In both membrane cases, the I-BAR domains have purely repulsive direct interactions with each other and are attracted to each other due to curvature coupling and Casimir-like forces mediated by the membrane as well as a membrane-composition-mediated force that occurs only as PIP2-like membrane beads cluster to the ends of the I-BAR domains (*26*). Thus, the functional difference between the PS-like membrane (i.e., 0% PIP2-like membrane) and the PIP2-like membrane is the additional attraction to the ends of the I-BAR domain for only a small percentage of the membrane beads. Consistent with our observation on GUVs, we found that I-BAR domain clustering occurs on both PS-like (i.e., 0% of PIP2-like membrane) and PIP2-like membranes (2% and 5% of PIP2-like membranes, Fig. 2D, Top). Moreover, we found that the addition of PIP2-like membrane beads increases the aggregation of I-BAR domains. Without PIP2-like membrane beads, we found a high probability of free or small aggregates containing less than five I-BAR domains. In the presence of PIP2-like membrane beads, there is a significant decrease in free or small aggregates and a corresponding increase of larger aggregates with five or more I-BAR domains (Fig. 2D Top and Fig. 2E). Importantly, it has been shown that I-BAR domains can induce stable PIP2 microdomains upon membrane binding (*32, 47*). Indeed, we found an enrichment of PIP2-like membrane beads near I-BAR domain aggregates (Fig. 2D, Bottom, blue dots). In the 2% PIP2-like membranes, we found 27% of neighboring membrane beads are PIP2-like, which is an approximate 14 times enrichment of PIP2-like membrane beads around the I-BAR domain aggregates compared to the total 2% concentration of PIP2-like beads on the membrane. Similarly, in the 5% PIP2-like membranes, we found 33% of neighboring membrane beads are PIP2-like, which is a 6.7 times enrichment of PIP2-like membrane beads to the aggregates compared to the total 5% concentration of PIP2-like membrane beads. The enrichment of PIP2-like membrane beads around I-BAR domain aggregates depletes PIP2 in the bulk membrane that is not adjacent to an I-BAR domain aggregate. In other words, the PIP2 percentages in the membrane adjacent to the I-BAR domain aggregates are enriched to 27% and 33% while the rest of the membrane is depleted to 0.7% and 3.4% of PIP2-like membrane beads for 2% and 5% PIP2-like membrane systems, respectively. Our results thus indicate positive feedback of the assembly of I-BAR domain aggregates mediated by PIP2 and the enrichment of PIP2 around the aggregates facilitate further I-BAR domain recruitment. Altogether, our simulation and experimental reconstitution results indicate the key role of PIP2 in IRSp53 clustering on membranes.

It was reported that in bulk, as well as on small vesicles containing PIP2, IRSp53 and VASP interact directly via their SH3 domain and PR domain, respectively (*15*). We thus examined this interaction in our reconstituted system. We purified full-length human VASP and labelled it with AX488 dyes (AX488-VASP). When incubating PIP2-GUVs with AX488-VASP (4 nM of VASP tetramer) together with unlabeled IRSp53 (16 nM of IRSp53 dimer), we observed that VASP is recruited on GUV membranes (Fig. 2F Top, arrows), which is not the case in the absence of IRSp53 (Fig. 2F, Bottom). This finding is consistent with the previous observations that IRSp53 and VASP interact directly in solution, on model membranes, and in filopodia (*15, 16, 18, 19*). Notably, we found clusters of VASP on GUV membranes (Fig. 2F, arrows) that are reminiscent of what has been observed in live cells (Fig. 1B and (*15*)). Furthermore, for GUVs having IRSp53-generated membrane tubules, we observed that VASP is recruited in these tubules (Fig. S3 and Movie S3). Our results indicate that IRSp53 can recruit VASP into clusters on flat membranes, and to negatively curved membranes, a characteristic of filopodial membranes.

### Self-assembly of IRSp53, VASP, fascin and actin on PIP2-membranes generates actin-filled membrane protrusions

We next assessed if IRSp53, VASP, actin and fascin can self-organize on PIP2-membranes to drive protrusion formation, as hypothesized above. We kept the bulk concentration of IRSp53 and VASP relatively low (16 nM of IRSp53 dimer and 4 nM of VASP tetramer) to allow the formation of IRSp53-VASP clusters on PIP2-membrane as seen in cells. To visualize actin, we used AX488 labelled globular actin (G-actin, ∼10% −27% AX488 labelled, total actin concentration 0.5 μM). To ensure that actin polymerization occurs at the membrane only, as in cells, and not in solution, we included capping protein (CP, 25 nM) and profilin (0.6 μM) in the protein mixture. Capping protein binds to the barbed ends of filamentous actin (F-actin) with high affinity (K_d_ = 0.1 nM) and inhibits F-actin elongation in the bulk (*1, 48*). In cells, the majority of G-actin is associated with profilin (K_d_ = 0.1 μM) (*49, 50*), which suppresses spontaneous actin nucleation in the bulk. Additionally, it was shown that at high ionic strengths, profilin is required for VASP to be more effective in actin polymerization (*51*). By performing pyrene actin polymerization assays, we verified that in our experimental conditions VASP increases actin polymerization in the presence of profilin, and CP suppresses F-actin elongation (Fig. S4, solid lines). Finally, to introduce actin filament bundling as in filopodia, we used fascin at 250 nM (*10*).

By incubating PIP2-GUVs with IRSp53, VASP, actin, fascin, CP and profilin, we observed inward membrane tubes filled with actin on the GUVs (Fig. 3A). The tubes are not static, but move rapidly inside GUVs (Movie S4 and S5). In addition to the tubes, there are actin signals on GUV membranes, indicating the formation of an actin shell on the membrane (Fig. 3A). On average, 32% of the GUVs had tubes, and 93% of these tube-positive GUVs had at least one tube filled with actin (N = 140 GUVs, n = 3 sample preparations, Fig. S5 A). To verify the presence of F-actin inside the tubes, we performed experiments using unlabeled G-actin and included AX488 phalloidin (0.66 μM) in the protein mixture, given that AX488 phalloidin binds to F-actin and its fluorescence is higher on F-actin than in the bulk (*52*). We verified by pyrene actin assays that the actin polymerization activity of VASP and the suppression of F-actin elongation by CP are preserved in the presence of phalloidin (Fig. S4, dashed lines). As shown in Fig. 3B, we observed clear AX488 phalloidin signals inside tubes as well as on GUV membranes, confirming the presence of F-actin (Movie S6 and S7). In the presence of phalloidin, on average, 55% of the GUVs had at least one actin-filled tube (N = 366 GUVs, n = 7 sample preparations, and Fig. S5 B). Using the fluorescence signals of AX488 phalloidin, we quantified that on average there are 11 actin filaments in the tubes (Fig. S5 C), comparable to what has been reported in cells (*9*). Besides tubes, we observed membrane deformations on the GUV membranes independent of the presence of phalloidin (Fig. S6, arrows). The deformed bulge shape of the GUV membranes is reminiscent of membrane deformation driven by the I-BAR domain of IRSp53 on GUVs (*53*), and indicates the presence of pushing forces acting on the GUV membranes towards the interior of the GUVs. Together, these results show that IRSp53, VASP, actin and fascin can spontaneously organize locally on PIP2-membranes to generate actin-based membrane protrusions.

**Fig. 3.**
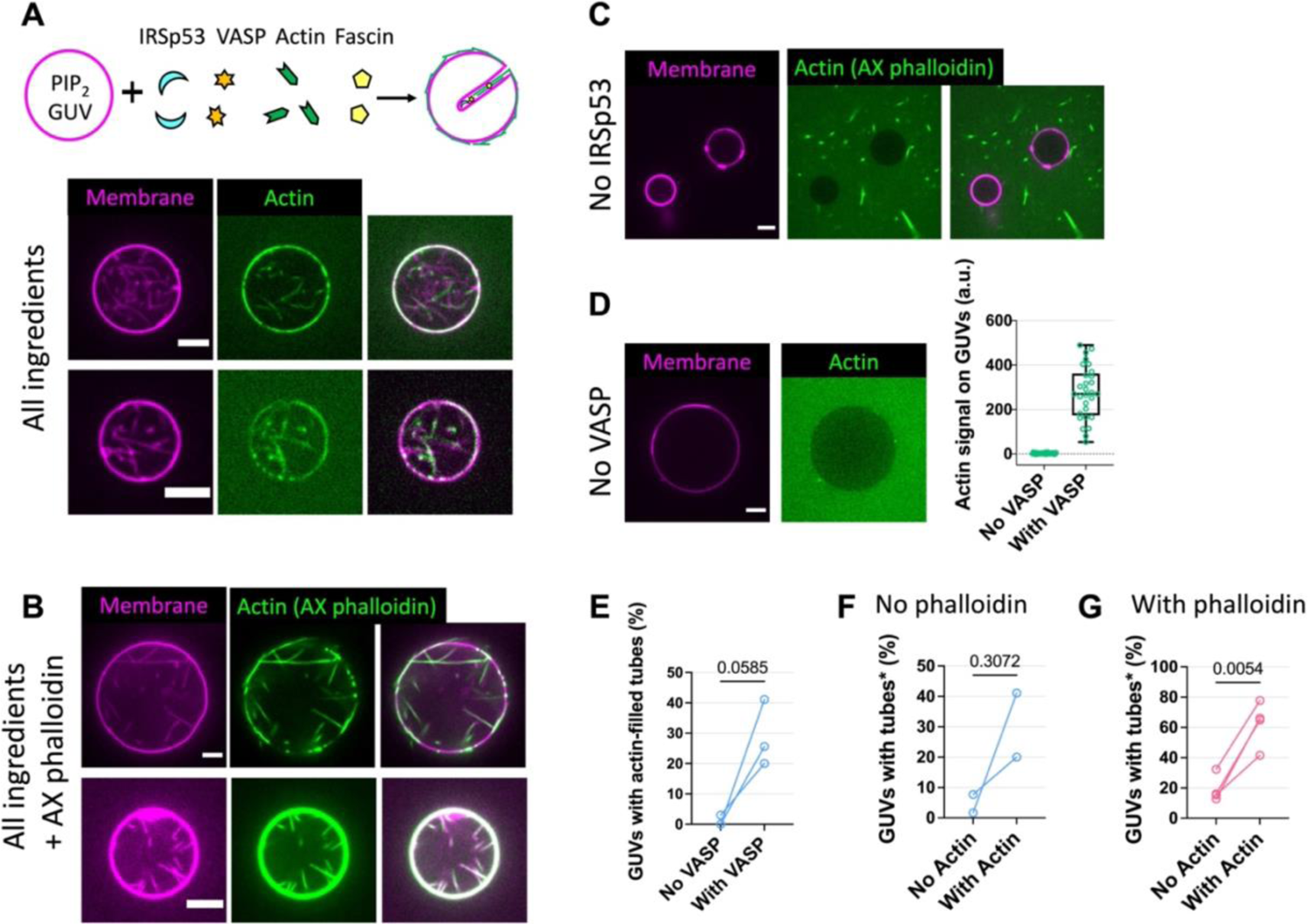
IRSp53 and VASP synergistically drive the formation of actin-filled membrane protrusions. (A) Top: cartoons depict GUVs incubated with IRSp53, VASP, actin and fascin (capping protein and profilin are not shown here), and the emergence of inward membrane tubes on the GUVs. Middle and Bottom: Two representative confocal images of GUVs incubated with IRSp53 (16 nM dimer), VASP (4 nM tetramer), actin (0.5 μM, 10% AX488 labelled, shown in green), fascin (250 nM), capping proteins (25 nM) and profilin (0.6 μM). Color codes: magenta, membranes and green, actin. (B) Two representative confocal images of GUVs in experimental condition identical as in (A) besides that AX488 phalloidin was included and there was no AX488 labelled actin in the protein mixture. (C) Identical experimental condition as in (A) but IRSp53 was absent. (D) Left: Representative confocal images of GUVs incubated with all the protein components as in (A) but no VASP, “No VASP”. Right: pixel-averaged fluorescence signals of actin on the contours of GUVs (GUV images were taken by focusing on the equatorial plan of the GUVs). N = 29 GUVs “No VASP” and N = 29 GUVs, “With VASP”, n = 1 sample preparation (please see Fig. S7A for the results of another two sample preparations). (E) Percentages of GUVs having actin-filled membrane protrusions in the absence (“No VASP”) and presence (“with VASP”) of VASP. GUV numbers, N = 42, 41, 33 for “No VASP” condition, and 56, 39, 45 for “With VASP”, n = 3 sample preparations. Statistical tests: (1) chi-squared test on data pooled from the 3 sample preparations, *p* < 0.0001, and (2) paired *t* test considering the 3 sample preparations individually, *p* = 0.0585. (F and G) Percentages of GUVs having tubes in the absence (“No Actin”) and presence (“With Actin”) of actin. *In the case of “With Actin”, GUVs were counted only when having actin-filled membrane tubes. (F) In the absence of phalloidin, “No phalloidin”, and used AX488 labelled actin. N = 39 and 60 for “No Actin” and N = 45 and 56 “With Actin”. n = 2 sample preparations. Statistical tests: (1) chi-squared test on data pooled from the 2 sample preparations, *p* < 0.0001, and (2) paired *t* test considering the 2 sample preparations individually, *p* = 0.3072. (G) In the presence of AX488 labelled phalloidin, and no AX488 labelled actin. GUV numbers, N = 31, 31, 26, 39 “No Actin”, and N = 54, 57, 41, 56 “With Actin”. n = 4 sample preparations. Statistical tests: (1) chi-squared test on data pooled from the 4 sample preparations, *p* < 0.0001 and (2) paired *t* test considering the individual 4 sample preparations, *p* = 0.0054. (A)-(D) Scale bars, 5 μm.

### IRSp53 is indispensable for protrusion formation by recruiting VASP to facilitate actin polymerization in protrusions

To reveal the contribution of individual protein components in the generation of actin-filled membrane protrusions, we performed loss-of-function assays by removing one protein component at a time. To ensure that protein activities are identical in the loss-of-function assays, we performed paired experiments in which we used the same batches of GUVs and protein stocks in each independent sample preparation. Then, in each sample preparation, we compared the efficiencies of the generation of actin-filled tubes in the absence of a protein of interest, and the reference condition where all the proteins are present. To this end, we counted the number of GUVs with and without tubes, and if an actin signal was readily detected in at least one or more tubes of the GUVs.

When IRSp53 was absent, we found none of the GUVs had tubes (Fig. 3C, N = 57 GUVs, n = 3 sample preparations, in the presence of AX488 phalloidin). Thus, in our experimental condition, IRSp53 is essential for the generation of actin-filled membrane tubes. In the absence of VASP, we observed a nearly complete lack of actin signal on the GUV membranes (Fig. 3D and Fig. S7A), and nearly no GUV had actin-filled tubes (out of a total of 116 GUVs, n = 3 sample preparations, only 8 GUVs were found with tubes, in which only 1 GUV had actin-filled tubes; Fig. 3E and Fig. S7B). Our observations thus indicate the key role of VASP in the generation of membrane tubes, via recruiting actin on GUVs. Given that VASP is an actin polymerase, we assessed if VASP facilitates actin polymerization in tubes by using AX488 phalloidin. Indeed, we observed that due to the presence of VASP, there is an increase in the number of GUVs having actin-filled tubes (Fig. S8, A and B) and an increase in the number of actin filaments in the tubes (Fig. S8 C). We note that in the presence of phalloidin, there are actin-filled tubes on GUVs even when VASP is absent (Fig. S8 D). This observation indicates that phalloidin aids actin polymerization even when CP and profilin are present. To understand how phalloidin influences actin polymerization in our system, we performed pyrene actin polymerization assays. We observed that phalloidin facilitates actin nucleation (Fig. S4, compare the two magenta lines), consistent with a previous report (*54*). It was shown in vitro that in the bulk, IRSp53 (after being activated by Eps8) and its I-BAR domain can interact with F-actin and induce actin bundle formation (*20, 24*). Thus, in the absence of VASP, phalloidin facilitates actin polymerization, and IRSp53 recruits F-actin to GUV membranes, resulting in the formation of actin-filled tubes. Consistently, we found that the percentages of GUVs having actin-filled tubes is higher in the presence of phalloidin compared to its absence (Fig. S5 D). Taken together, our results show that through IRSp53-driven recruitment to PIP2-membranes, VASP plays a key role in protrusion generation via its actin polymerization activity.

Biophysical studies using reconstitution systems and theoretical modelling have revealed how the interplay between the mechanical properties of membranes and actin bundles/networks determines the formation of actin-driven membrane protrusions (*11, 55, 56*). Given that actin’s role in the initiation of IRSp53-driven cellular protrusions is not fully understood (*17*), we assessed if actin facilitates IRSp53-based protrusion formation using our reconstitution systems. In the absence of actin, we observed 2% to 34% of GUVs having tubes (Fig. 3 F and G, regardless of phalloidin). These tubes were generated by IRSp53 since when IRSp53 is absent, no GUV has tubes (Fig. 3C). Notably, we observed a global increase in the amount of GUVs having tubes due to the presence of actin (Fig. 3 F and G). This effect is more pronounced when phalloidin is present (Fig. 3G, N = 127 and 208 total GUVs, without and with actin, respectively. n = 4 sample preparations). In the absence of phalloidin, in a sample preparation where there was a relatively high amount of GUVs with tubes in the absence of actin (34.1%, N = 41 GUVs), the addition of actin did not aid tube generation (25.6%, N = 39 GUVs). However, in experimental sets where the amount of GUVs with tubes was low (less than 10%) in the absence of actin, the addition of actin increased the amount of GUVs with tubes (Fig. 3F, N = 99 and 101 total GUVs, without and with actin, respectively, n = 2 sample preparation). Together, these results indicate that actin contributes to IRSp53-based tube generation. Furthermore, given that phalloidin facilitates actin nucleation on membranes in our system, our results point out the essential role of actin nucleation to enhance actin’s function in protrusion formation in cells.

### Fascin is not required for protrusion generation but enhances protrusion elongation and stability

Given that fascin is the specific actin bundler in filopodia, we assessed its role in protrusion formation. In the presence and absence of fascin, we did not observe significant differences considering the number of GUVs having actin-filled tubes, regardless of the absence (Fig. 4A and Fig. S9 A) or presence of phalloidin (Fig. S9 B and C). Furthermore, there is no significant difference in the amount of F-actin in tubes in the presence and absence of fascin (Fig. S9 D). Due to the rapid movement of tubes inside GUVs, we could not characterize the dynamics of tube generation and elongation. We thus assessed how fascin affects protrusion dynamics in live cells. To this end, we performed experiments using Rat2 cells that expressed GFP-tagged IRSp53 and mCherry-tagged fascin (Fig. 4B). We observed that the recruitment of fascin in IRSp53-based protrusions coincides with their elongation (Fig. 4 C and D). By tracking IRSp53-based protrusions, we also found that fascin significantly increases the growth rates of the protrusions (Fig. 4D), and that its over-expression decreases the frequency of their retraction (Fig. 4E). Thus, our in vitro reconstitution data demonstrate that actin polymerization facilitates the formation of IRSp53-dependent membrane protrusions, while the cell data suggest that fascin increases the growth rate and enhances the stability of these protrusions.

**Fig. 4.**
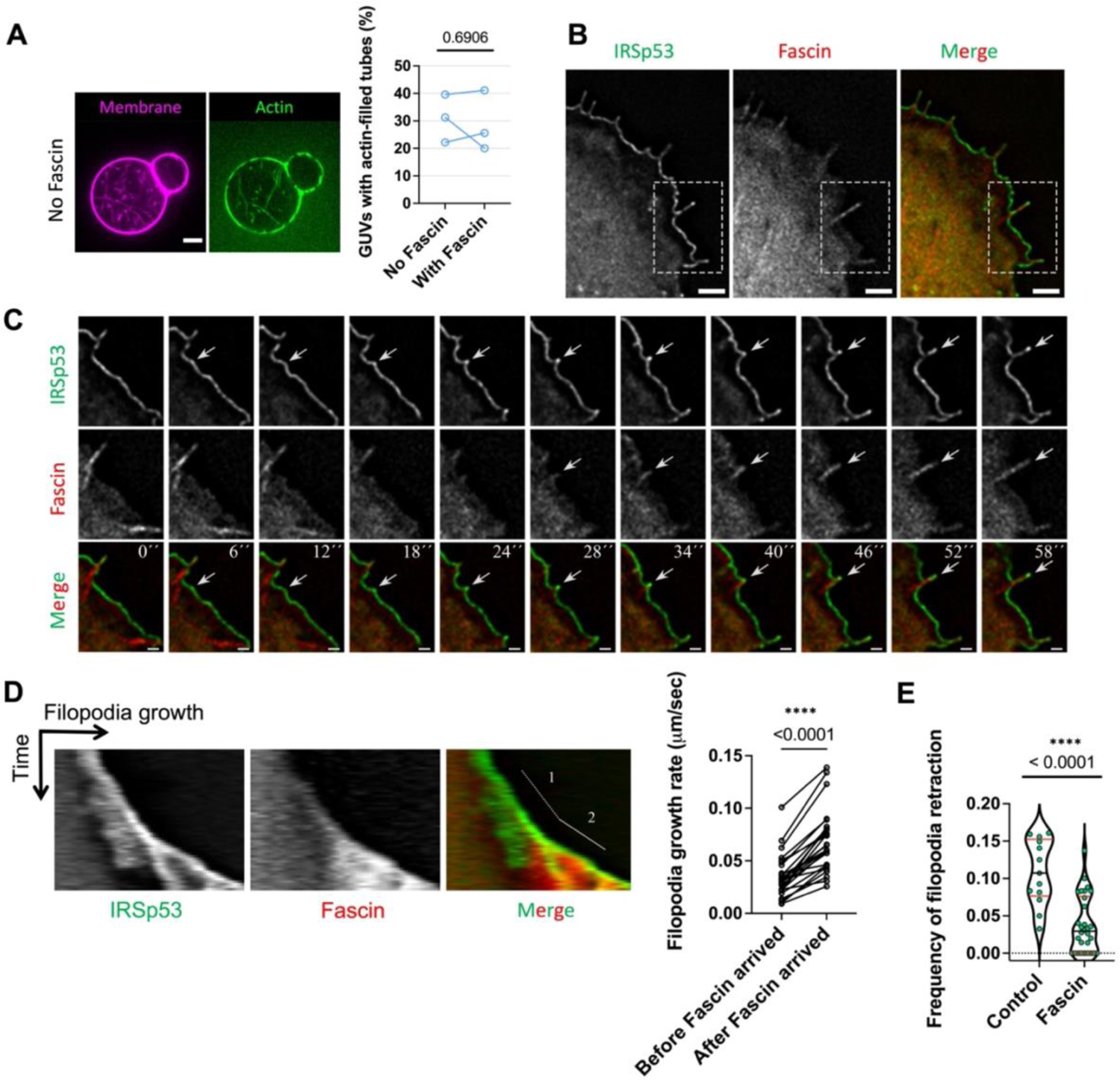
Fascin facilitates filopodia growth and prevents filopodia retraction. (A) Left: Representative confocal images of GUVs incubated with all the protein ingredients (IRSp53, VASP, actin, capping protein and profilin) and in the absence of Fascin. Right: Percentages of GUVs having actin-filled membrane tubes in the absence (“No Fascin”) and the presence (“With Fascin”) of Fascin. GUV numbers: N = 53, 32, 54 “No Fascin” and 56, 45, 39 “With Fascin”. n = 3 sample preparations. Statistical tests: (1) chi-squared test on data pooled from the 3 sample preparations, *p* = 0.8652, (2) paired *t* test considering the 3 sample preparations individually, *p* = 0.6906 (shown in the figure). (B) Widefield fluorescence image of a representative Rat2 cell transfected with pEGFP-N1-IRSp53L (IRSp53-GFP) and fascin-pmCherry (fascin-Cherry). Scale bar, 2 µm. (C) Time-lapse images of a single filopodia formation. Magnification of the indicated area shown in (B), white boxes. The white arrow indicates the appearance of IRSp53 clusters followed by the recruitment of fascin in filopodia formation. Scale bars, 2 µm. Time in sec. (D) *Left* panel: Representative kymograph of IRSp53 and fascin fluorescence signals on growing filopodia from Rat2 cell transfected with pEGFP-N1-IRSp53L (IRSp53-GFP) and fascin-pmCherry (fascin-Cherry). Number 1 and 2 in the kymograph indicate filopodia growth before (Number 1) and after (Number 2) the presence of fascin inside filopodia. *Right* panel: Quantification of filopodial growth rate before (“Before Fascin arrived”) and after (“After Fascin arrived”) the emergence of fascin inside the filopodia. N = 26 filopodia. Statistical test: Mann-Whitney nonparametric test, *p* = 0.00001633. (E) Frequency of filopodia retractions in Rat2 cells transfected with pEGFP-N1-IRSp53L (IRSp53-GFP) and with either empty-pmCherry (“Control”) or fascin-pmCherry (“Fascin”). The frequency of each event was calculated for the period of filopodia growth as the number of retractions per sec. The numbers of filopodia analysed were n=13 (“Control”) and n=30 (“Fascin”). Statistical test: Mann-Whitney nonparametric test, *p* = 0.000083802.

### IRSp53 regulation in cells revealed by membrane nanotube pulling experiments

Our results obtained using the reconstituted system demonstrate the function of IRSp53 in the generation of actin-filled membrane protrusions and show that IRSp53 is primed to self-assemble into clusters that drive the association of downstream partners (e.g., VASP, fascin) for protrusion growth. However, it remains unclear how cells regulate the activity of IRSp53 to control for instance protrusion formation at specific membrane locations. Indeed, with the abilities of IRSp53 to sense membrane curvature and to trigger actin assembly, we expect that any local deformation of the plasma membrane could induce filopodial growth. To reveal IRSp53’s possible regulation directly in cellulo, we generated artificial protrusions having identical topologies to filopodia by pulling membrane nanotubes (i.e., tethers) from the plasma membrane of Rat2 fibroblasts using optically trapped polystyrene beads and micromanipulation (Fig. 5A). The pulled nanotubes serve as tractable models to directly assess the recruitment of IRSp53 into tubular geometries and the possible actin assembly inside these nanotubes as a consequence of IRSp53 enrichment. Cells were transfected with either IRSp53’s I-BAR domain (I-BAR-eGFP), its putative membrane deforming and curvature sensitive region, or the full-length IRSp53 protein (IRSp53 FL-eGFP). The plasma membrane was exogenously labelled using the lipophilic Cell Mask^TM^ Deep Red stain. Rat2 I-BAR-eGFP expressing cells show strong I-BAR-eGFP fluorescence in pulled nanotubes (Fig. 5B) that was visible after nanotube formation (Fig. S10 and Movie S8), suggesting an innate preference of the I-BAR domain to sort into these nanotubes. To quantify the enrichment of proteins in the pulled nanotubes, we calculated sorting (*S*) heat map images from the membrane and protein fluorescence channels, and then determined the mean protein sorting value for a given nanotube (*S*_avg_) by averaging the maximum sorting value at each pixel position along the nanotube axis (Fig. 5C; see Materials and Methods for further details). Note that *S*_avg_ values greater than 1 indicate preferential protein enrichment for a given nanotube. We measured the *S*_avg_ of the I-BAR domain for multiple nanotubes (N = 19 nanotubes) and determined an ensemble average of 4.5 (Fig. 5D). This value is similar to previously reported sorting values obtained from nanotubes pulled from GUVs encapsulating the purified I-BAR domain (*33*).

**Fig. 5.**
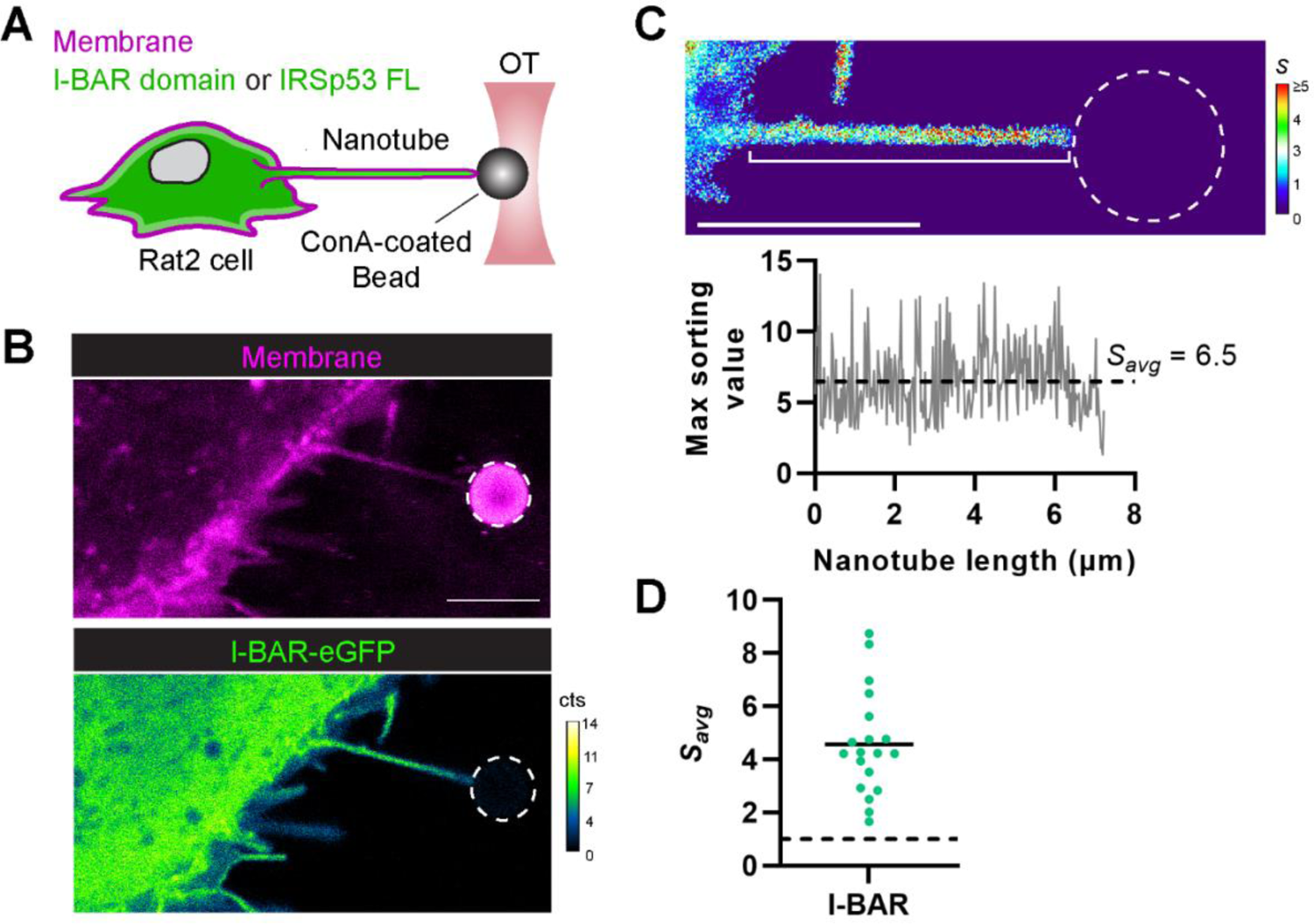
IRSp53’s I-BAR domain is robustly recruited into pulled membrane nanotubes. (A) Experimental setup for pulling membrane nanotubes using a concanavalin A (con-A)-coated bead trapped in an optical tweezer (OT). Rat2 fibroblasts, expressing eGFP fusions (green) of either IRSp53’s I-BAR domain or the full length (FL) IRSp53 protein, were labelled with Cell Mask^TM^ Deep Red plasma membrane stain (magenta); protein enrichment in the membrane nanotube was monitored by confocal fluorescence microscopy using single-photon avalanche detectors (cts, counts). (B) Representative confocal image of a pulled membrane nanotube from a Rat2 cell expressing IRSp53’s I-BAR domain showing high enrichment of the I-BAR domain. (C) Top: Calculated sorting map of the nanotube in (B) with low sorting (*S*) values in blue and high *S* values in red. Bottom: Plot of the maximum sorting value at each pixel position along the length of the nanotube (white bracket in the sorting map) and the mean sorting value for the protein (*S*_avg_). (D) Measured *S*_avg_ values for IRSp53’s I-BAR domain in pulled nanotubes (N = 19 nanotubes). *S*_avg_ > 1 (dashed black line) indicate protein enrichment. Black solid line, mean of the data points. Dashed white circles in the figure outline the trapped bead. Scale bars, 5 µm.

Contrary to the robust enrichment of the I-BAR domain, we observed a more complex behavior for the sorting of the full-length IRSp53 protein in nanotubes that is dependent upon the local cellular membrane activity near the sites of the nanotubes. Pulled nanotubes from IRSp53 FL-eGFP expressing cells show strong IRSp53 signal only when the nanotube was pulled near zones exhibiting “active” processes of membrane remodeling (Fig. 6A), such as lamellipodia and membrane ruffles, which were frequently found at the cell leading edge. However, no IRSp53 FL-eGFP fluorescence was observed in nanotubes when pulled from “non-active” zones (Fig. 6B), such as those found near the cell trailing edge. Subsequent imaging over time showed that nanotubes pulled from “active” and “non-active” regions stably retained either the presence or absence of IRSp53, respectively (Fig. S11). As seen in Fig. 6C, nanotube *S*_avg_ values for IRSp53 FL near “active zones” (N = 13 nanotubes) averaged 2.8 and was comparable to the I-BAR domain case (Fig. 5D). However, *S*_avg_ values for IRSp53 FL near “non-active zones” (N = 20 nanotubes) averaged 0.2, indicating an exclusion of the protein from the nanotubes. The dichotomy of sorting behaviors for IRSp53 FL is completely opposite to the results we observed for the I-BAR domain, where the I-BAR domain sorting is consistently stronger and did not depend on the cellular region where the nanotube pulling was performed. Together, our results highlight that IRSp53’s curvature sensing ability is innately part of the I-BAR domain (*33, 34*) and further suggest that cellular regulation mechanisms are at play in modulating IRSp53’s activity.

**Fig. 6.**
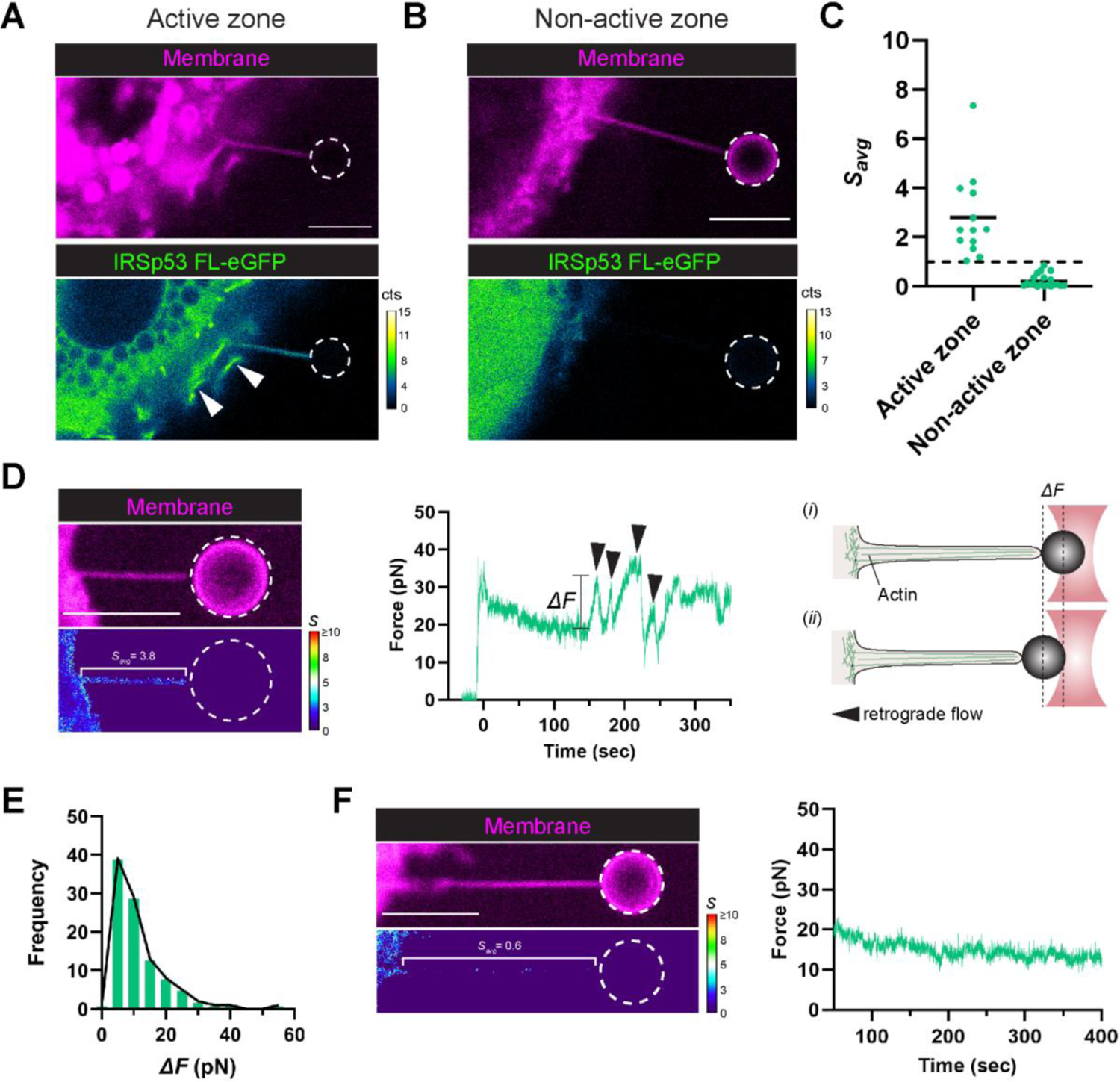
IRSp53 is recruited into membrane nanotubes pulled from highly active cellular zones and coincides with actin development. (A) and (B) Representative confocal images of pulled membrane nanotubes from Rat2 cells expressing the full-length IRSp53 protein. (A) Pulled nanotubes near cellular zones of active membrane remodeling, such as membrane ruffling (white arrowheads), exhibit enrichment of IRSp53 in the nanotube. (B) Pulled nanotubes near non-active zones instead show no enrichment of IRSp53. (C) *S*_avg_ values for IRSp53 were determined from active and non-active zones. Dashed black line, *S*_avg_ = 1. Black solid lines, mean of the data points. Active zone, N = 13 nanotubes; Non-active zone, N = 20 nanotubes. (D) IRSp53 recruitment corresponds with eventual actin development within the pulled nanotube. Representative sorting map (left) and the corresponding force plot (center) for a nanotube showing high IRSp53 recruitment. Peaks in the force plot (black arrowheads) are signatures of actin in the tube, and arise when retrograde flows outcompete actin polymerization (at the nanotube tip) causing bead displacement towards the cell body and hence a rise in the force (right, *i* and *ii*). (E) Distribution of the force peak magnitudes (*ΔF*). Sample size, 100 peaks. (F) Representative sorting map (left) and the corresponding force plot (right) for a nanotube showing no IRSp53 recruitment and hence no actin development. Color maps: low sorting (*S*) values in blue, and high *S* values in red. Dashed white circles in the figure outline the trapped bead. Scale bars, 5 µm.

We observed that the sorting of IRSp53 FL has a direct consequence on the assembly of F-actin within the nanotubes. Nanotubes with high IRSp53 sorting (Fig. 6D) exhibited characteristic signatures of F-actin in the measured force profile and behaviors mimicking those of true filopodia. The observed bead displacement within the optical trap (i.e., a rise in the force, ΔF) indicates that F-actin development has reached the end of the nanotube, allowing retrograde and contractile forces to transmit from the cell body to the bead. The magnitude of the force peaks generates traction forces of 5−10 pN (Fig. 6E) and are of the same order of magnitude as those reported for true filopodia (*57, 58*). Additionally, events of nanotube curling (i.e., helical buckling) were also frequently observed (Fig. S12), as previously shown for filopodia (*58*). In essence, IRSp53 FL positive nanotubes eventually mature into a “pseudo” filopodium. However, nanotubes with no IRSp53 sorting (Fig. 6F) exhibit a constant force profile and consequently no F-actin development was observed. Together, our tube pulling results show that though IRSp53 is curvature sensitive through its I-BAR domain, its activity is tightly controlled within cells to constrain local F-actin development for protrusion generation.

## Discussion

How cells control the formation of protrusions such as filopodia and microvilli at specific membrane locations remains largely unclear. Using in vitro reconstitution systems, we demonstrated for the first time that a minimum set of proteins comprised of the membrane curvature sensor IRSp53, the actin polymerase VASP, and actin can spontaneously organize to generate actin-based membrane protrusions. By performing loss-of-function assays, we investigated the function of proteins in protrusion formation. We demonstrated that IRSp53 is the essential player in protrusion generation by recruiting VASP to promote local actin assembly. Thus, by using a precisely controlled reconstitution system, our study provides a strong support to the previously proposed mechanism of IRSp53-VASP-driven filopodia generation that was largely based on cell biology studies (*15, 16, 18*).

A key finding of our in vitro assay is that purified full-length IRSp53 is active in the absence of its activators such as Cdc42. Our results indicate that once IRSp53 binds to the plasma membrane, it can self-assemble into clusters and readily recruit VASP to generate actin-filled protrusions. Indeed, it was reported that when expressing a non-regulated constitutively active mutant of IRSp53 in cells, an aberrant explosive formation of filopodia was observed (*21*). Thus, to prevent generating unwanted protrusions, cells must have a tight regulation of IRSp53, i.e., how much, when, and where to activate IRSp53. To reveal the regulation of IRSp53 activity in cells, we generated filopodia-like membrane geometry in live cells by pulling membrane nanotubes from the plasma membrane. We discovered that IRSp53 can only be recruited to nanotubes pulled from active membrane regions of the cell where the membrane exhibits dynamic ruffles, but not from regions where no membrane remodeling was observed. Moreover, actin polymerization extending throughout the IRSp53-enriched nanotube was detected, creating in essence a “pseudo” filopodium. Our finding is consistent with the location of IRSp53’s activator Cdc42 at the front edge of cells where ruffling and protrusions develop (*59, 60*), and with IRSp53’s function in inducing Rac-dependent membrane ruffling (*27, 30*). Importantly, our results indicate that to finely regulate IRSp53 activity, it is probably even more important to keep IRSp53 inhibited such that it is away from membranes. Indeed, it was shown that the binding of 14-3-3 to IRSp53 counteracts the activation by Cdc42 and other downstream cytoskeleton effectors (*21*). Notably, 14-3-3 binding to phosphorylated IRSp53 keeps it in an inhibited state, resulting in impaired filopodia formation and dynamics (*21, 30, 31*). Collectively, our results reflect a finely tuned regulatory mechanism where IRSp53 is kept in its inhibition state by phosphorylation and binding to 14-3-3, and activated by binding to activators such as Cdc42. Controlling IRSp53 by three different signaling pathways, phosphorylation, Cdc42 binding and PIP2 binding thus ensures its precise spatial-temporal regulation such that IRSp53 is activated only at specific regions of cells.

It was shown previously that PIP2-binding by IRSp53’s I-BAR domain is required for the generation of plasma membrane protrusions (*26*). Indeed, our in vitro work and coarse-grained simulation results showed that PIP2 is key for the assembly of these IRSp53 clusters: higher PIP2 concentration results in larger I-BAR domain clusters. Importantly, our simulation results showed an at least 7-fold enrichment of PIP2 around the I-BAR domain clusters. Our finding is consistent with the previous reports of BAR and I-BAR domains inducing local and stable PIP2 clusters (*32, 47, 61*). Notably, it was proposed that these PIP2 clusters are transiently interacting with BAR domains such that PIP2 in the clusters is available for the recruitment of downstream partners having PIP2 binding motifs (*32, 47, 61*). We thus propose that besides specific protein-protein interactions, IRSp53-induced PIP2 clusters could facilitate not only the further recruitment of IRSp53 but also other proteins such as actin nucleation promoting factors, actin polymerases and actin-binding proteins for protrusion formation and regulation (*16, 28, 62-65*).

We observed that clusters of IRSp53 preceding those of VASP on the plasma membrane of Rat2 cells at the onset of filopodia formation, in agreement with what reported by other studies using MEF cells and COS-7 cells (*15, 21*). VASP clustering has been shown to be an important factor to increase the processivity of VASP for actin filament elongation (*35, 39*). Consistently, we showed that VASP clustering via IRSp53 efficiently polymerizes actin on PIP2-membranes and in protrusions. Moreover, we observed an increased number of actin filaments in protrusions when VASP is present. Of note, a recent study using B16F1 cells showed that VASP clustering for filopodia formation is initiated by lamellipodin, but not by IRSp53 (*42*). This study and our work thus suggest that VASP requires membrane-interacting partners to bring it to membranes and induce its clustering to drive outward membrane deformation. The differences in the mechanisms of recruiting VASP on membranes maybe cell-type dependent, i.e., cells may utilize different pathways to induce membrane deformation via VASP. Future work is required to decipher how and under which physiological conditions cells use different precursor proteins to cluster VASP on the plasma membrane for the formation of membrane protrusions.

In cells, filopodia formation involves multiple protein complexes (*66*). Our in vitro work provides direct evidence that actin filament assembly facilitates IRSp53-based protrusion generation. This is consistent with what was proposed previously based on EM and cell biology experiments (*17*). In our assay, actin assembly is driven by VASP, given that when VASP was eliminated, we observed nearly no actin signal on GUVs (Fig. 3D). We anticipate that other actin assembly factors can play similar roles as VASP. Indeed, it was reported that IRSp53 interacts directly with formins, such as mDia1, in filopodia formation (*28*). Moreover, actin nucleation promoting factors WAVE2 and N-WASP were shown to synergize with IRSp53 to generate filopodia (*16, 28*). In our in vitro system, fascin is not essential for protrusion initiation. However, by observing the dynamics of filopodia in Rat2 cells, we found that fascin enhances the elongation and stability of filopodia. Our result thus supports the notion that fascin mechanically strengthens filopodia by bundling actin filaments together, thus facilitating filopodial extension and stabilization (*4–7*). Notably, Eps8, another actin bundler, has been reported to form a complex with IRSp53 during filopodia formation (*18, 20*). Besides bundling, Eps8 can cap the barbed end of actin filaments. The dual function of Eps8 on actin filaments is fine-tuned in cells, allowing cells to generate filopodia via different signaling networks depending on cellular contexts (*18*). To understand how the abovementioned proteins regulate dynamic filopodia formation, elongation, retraction, and force generation, future work is required to elucidate the interplay between these proteins in the context of protrusion formation.

Altogether, our work demonstrates that IRSp53 is an efficient protrusion initiator: once being activated, IRSp53 readily triggers the cascade of protrusion formation. Thus, to avoid the uncontrolled formation of protrusions, it is critical for cells to have a strict and precise control on when and where to activate or unlock IRSp53 from its inhibitory state

## Materials and Methods

### Pyrene actin polymerization assay

Polymerization assays were based on measuring the fluorescence change of pyrenyl-labeled G-actin (λexc=365 nm, λem=407 nm). Experiments were carried out on a Safas Xenius spectrofluorimeter (Safas, Monaco). Polymerization assays were performed in buffer containing 60 mM NaCl, 1 mM MgCl_2_, 0.2 mM EGTA, 0.2 mM ATP, 10 mM DTT, 1 mM DABCO, 5 mM Tris pH 7.5 and 0.01% NaN_3_ in the presence of actin (2 μM, 5% pyrenyl labeled), profilin (2.4 μM), VASP (15 nM in tetramer), CP (25 nM) and phalloidin (2 μM).

### GUV experiments

#### Reagents

Brain total lipid extract (TBX, 131101P), brain L-α-phosphatidylinositol-4,5-bisphosphate (PIP_2_, 840046P) were purchased from Avanti Polar Lipids. BODIPY-TR-C5-ceramide, (BODIPY TR ceramide, D7540) was purchased from Invitrogen. Alexa Fluor 488 tagged phalloidin (AX488 phalloidin) was purchased from Interchim. β-casein from bovine milk (>98% pure, C6905) and other reagents were purchased from Sigma-Aldrich.

#### GUV preparation

The lipid mixture used contain total brain extract supplemented with 5 mol% PI(4, 5)P2 at 0.5 mg/mL in chloroform. If needed, the lipid mixture is further supplemented with 0.5 mol% BODIPY TR ceramide.

GUVs were prepared by using the polyvinyl alcohol (PVA) gel-assisted vesicle formation method as previously described (*67*). Briefly, a PVA gel solution (5 %, w/w, dissolved in 280 mM sucrose and 20 mM Tris, pH 7.5) warmed up to 50 °C was spread on clean coverslips (20mm × 20mm). The coverslips were cleaned by ethanol and then ddH2O twice. The PVA-coated coverslips were incubated at 50 °C for 30 min. Then, around 5 μl of the lipid mixture were spread on the PVA-coated coverslips, followed by placing them under vacuum at room temperature for 30 min. The coverslips were then placed in a petri dish and around 500 μl of the inner buffer was pipetted on the top of the coverslips. The inner buffer contains 50 mM NaCl, 20 mM sucrose, 20 mM Tris pH 7.5. The coverslips were kept at room temperature for at least 45 min, allowing GUVs to grow. Once done, we gently “ticked” the bottom of the petri dish to detach GUVs from the PVA gel. The GUVs were collected using a 1 ml pipette tip with its tip cut to prevent breaking the GUVs.

### Protein purification and labelling

Muscle actin was purified from rabbit muscle and isolated in monomeric form in G-buffer (5 mM Tris-Cl^-^, pH 7.8, 0.1 mM CaCl_2_, 0.2 mM ATP, 1 mM DTT, 0.01% NaN_3_) as previously described (*68*). Actin was labeled with Alexa 488 succimidyl ester-NHS (*69*).

The genes encoding human IRSp53 and VASP were provided by Prof. Roberto Dominguez (University of Pennsylvania), and Prof. Jan Faix (Hannover Medical School), respectively, and sub-cloned into the pGEX-6P-1 vector (Cytiva). The expression plasmids were transformed into BL21 (DE3) competent cells. A single colony was resuspended in 10 mL LB medium containing 100 μg/mL ampicillin, and cultured at 37 °C overnight. Then, the starter culture was inoculated into 1 L LB medium containing 100 μg/mL ampicillin, and cultured at 37 °C until OD600 reached 0.6. The protein expression was induced with 0.2 mM isopropyl-D-1-thiogalactopyranoside (IPTG) overnight at 15°C. After harvesting the cells, the pellets were resuspended and sonicated in a buffer containing 20 mM Tris-HCl (pH 7.4), 500 mM NaCl, 1 mM EDTA, 1 mg/mL lysozyme, 1% Triton X-100, 1 mM PMSF, and 1 mM DTT, followed by affinity purification with GSTrap FF (Cytiva). The GST-tag was removed with PreScission protease (Cytiva). The proteins were concentrated to 50−100 µM using the Amicon Ultra centrifugal filters (Merck). For protein labelling, the Alexa Fluor maleimide (Thermo Fisher Scientific) was added to the protein solution, and the reaction was allowed to proceed at 4 °C overnight in the dark to protect it from light. The labelled proteins were further purified and separated from free Alexa Fluor dye using a Superdex 75 10/300 GL gel filtration column (Cytiva) using either an ÄKTA protein purification system (Cytiva) or NGC Chromatography System (Bio-Rad). The proteins were concentrated using the Amicon Ultra centrifugal filters (Merck) by replacing the buffer with 20 mM Tris-HCl (pH 7.4), 150 mM NaCl, and 1 mM DTT, then 0.1 % (w/v) methylcellulose was added. The purified proteins were frozen in liquid nitrogen and stored at −80 °C before use. The labelling efficiency of Alexa 488 IRSp53 (dimer) is 0.94, and that of Alexa 488 VASP (tetramers) is 3.0. Recombinant human profilin I was expressed in BL21 (DE3) Star and purified as described (*70*). The plasmid for the expression of mouse capping protein α1β1 was cloned in a pRSFDuet-1 vector, providing a 6xHis tag at the N terminus of the α1 subunit. Mouse capping protein α1β1 was expressed in BL21 (DE3), and was purified as previously described (*71*).

The plasmid expressing GST-HsFascin1 was provided by Prof. David Kovar (University of Chicago). The homo sapience Fascin1 full length protein (UniprotID: Q16658) was produced in BL21(DE3) bacterial cells from the pGEX-4T-3 vector as described previously with some modifications (*10*). In brief, the bacterial cells were grown for 24 hours at RT (around 20C) in auto-induction media (AIMLB0210, Formedium, UK), inoculated with overnight grown starter culture. The cells were harvested by centrifugation and resuspended in PBS buffer supplemented with 1 mM DTT, 10 mM CaCl2, cOmplete™ Protease Inhibitor Cocktail (#11697498001, Merck), and DNAseI (Sigma, DN25-1G). The cells were disrupted by sonication and soluble fraction after 1 hour centrifugation at 18 000 x g was applied to the Glutathione Sepharose 4 Fast Flow (#17513201, Cytiva). After 2 hours incubation at +4C, the beads were extensively washed with the same buffer and thrombin protease were added to beads, mixed well, and incubated over night at +4C. The next day the cleaved soluble fraction from the beads was collected and subjected to HiLoad 16/600 Superdex 75 gel filtration chromatography column, preequilibrated with 20 mM HEPES pH 8.0, 50 mM NaCl, 0.01% NaN3. The elution fractions containing the protein were collected, concentrated with 30 kDa MWCO VivaSpin column, flash-frozen in liquid nitrogen, and stored at −80C until experiments.

### GUV sample preparation and observation

For all experiments, coverslips were passivated with a β-casein solution at a concentration of 5 g.L^-1^ for at least 5 min at room temperature. Experimental chambers were assembled by placing a silicon open chamber on a coverslip.

Actin polymerization buffer contains 60 mM NaCl, 1 mM MgCl_2_, 0.2 mM EGTA, 0.2 mM ATP, 10 mM DTT, 1 mM DABCO, 5 mM Tris pH 7.5. GUVs were mixed sequentially with the ingredients, if present, in the following order: actin polymerization buffer, IRSp53, VASP, fascin, profilin, capping protein, phalloidin, GUVs and finally actin. The GUV-protein mixture was then pipetted using a pipette tip with its tip cut into the experimental chamber, followed by incubating at room temperature for at least 15 min before observation. The final concentration of proteins, if present, were: 16 nM IRSp53, 4 nM VASP, 250 nM fascin, 0.6 μM profilin, 25 nM capping protein and 0.5 μM actin. If needed, we diluted protein stocks in actin polymerization buffer such that the final concentrations of salt and ATP in the GUV-protein mixture maintained approximately constant. To visualize actin by confocal microscopy, depending on the experiments, we used either actin monomers having 10% − 27% fluorescently labelled with AX488, or unlabeled actin but included AX488 conjugated phalloidin (AX488 phalloidin) at a final concentration of 0.66 μM.

Single actin filaments were prepared by mixing actin monomers with the actin polymerization to reach a final actin concentration at 0.5 μM.

Samples were observed using either spinning disk confocal microscopes or a laser scanning confocal microscope. The two spinning disk confocal microscopes used are (1) Nikon eclipse Ti-E equipped with a Yokogawa CSU-X1 confocal head, a 100X CFI Plan Apo VC objective and a CMOS camera, (Prime 95B, Photometrics) and (2) Nikon eclipse Ti-E equipped with a 100x/1,4 OIL DIC N2 PL APO VC objective and an EMCCD camera (Evolve). The laser scanning confocal microscope is a Nikon TE2000 microscope equipped with an C1 confocal system and a 60X water immersion objective (Nikon, CFI Plan Apo IR 60XWI ON 1,27 DT 0,17).

### Quantification of AX488 actin on GUV membranes

The pixel-averaged AX488 actin fluorescence intensities on GUV membranes were obtained using a previously described method (*72*). Membrane fluorescence signals of GUVs were used to detect the contours of the GUVs by using the “Fit Circle” function in Fiji. Then, a 5 pixel wide band centred on the GUV contours were used to obtain the actin intensity profile of the band where the x-axis of the profile is the length of the band and the y-axis is the averaged pixel intensity along the band width. Actin intensity was then obtained by calculating the mean value of the intensity values of the intensity profile, following by subtracting the background intensity. To obtain the background intensity of AX488 fluorescence, we first manually drew a 5 pixel width line perpendicularly across the GUV membranes, and then, the background intensity was obtained by calculating the mean value of the sum of the first 10 intensity values and the last 10 intensity values of the background intensity profile, in which the x-axis of the profile is the length of the line and the y-axis is the averaged pixel intensity along the width of the line.

### Estimation of the number of actin filaments in protrusions

To estimate the number of actin filaments inside protrusions, we extracted fluorescence signals of actin in protrusions and normalized them by the fluorescence signals of single actin filaments in bulk prepared by using the same actin stock as those in the GUV-protein mixture. The microscope setting for image acquisition was identical for the GUV sample and for the corresponding single actin filaments in bulk. We performed the following steps to extract actin signals in protrusions and in bulk. We manually defined the ROI, a line with a width of 5 pixels drawn perpendicularly across protrusions or single actin filaments. We then obtained the actin fluorescent intensity profile of the line where the x-axis of the profile is the length of the line, and the y-axis is the averaged pixel intensity along the width of the line. The background intensity was obtained by calculating the mean value of the sum of the first five intensity values and the last five intensity values of the intensity profile. Finally, the actin fluorescence intensities were obtained by subtracting the background intensity from the maximum intensity value in the intensity profile. This image process was performed by using Fiji (*73*).

### Characterization of IRSp53 clustering on GUVs

To define clusters of IRSp53 on a surface of a GUV we segmented fluorescence images of the protein using a custom-made Fiji script based on the Rényi Entropy algorithm (*74*). We first created a mask of the fluorescent image in the IRSp53 channel using a Rényi Entropy threshold, then used the “analyse particles” function in Fiji to define clusters as a set of non-connected areas that have non-zero values in the mask image, permitting quantification of the number of clusters and their areas.

## Statistical analysis

For GUV experiments, all graphs and statistical analyses were performed using GraphPad Prism software version 9.3.1 for Mac (GraphPad Software, San Diego, CA, USA).

### Coarse-grained Model and Clustering Analysis

The coarse-grained (CG) model used here has been discussed in detail previously (*75, 76*). The low-resolution, phenomenological CG model contains a 3-bead quasi-monolayer membrane model and a curved I-BAR domain model. The quasi-monolayer model is highly tunable and can accommodate significant remodeling, which makes it appealing for the application to I-BAR domains. The 3-beads interact internally with two harmonic bonds with force constant of 25 *k_B_T* and equilibrium distance of 0.9 nm and a harmonic angle potential with force constant of 10 *k_B_T* and equilibrium angle of 180 degrees. The intermolecular forces using a soft pair potential, shown below, where *A* and *B* dictate the softness of the repulsion and the depth of the attraction and *r_0_* and *r_c_* are the onset of repulsion cutoff and attraction cutoff, respectively.

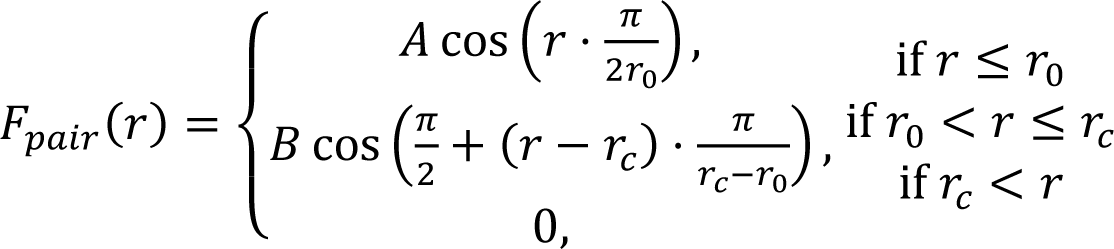

Both the membrane and the protein have an A value of 25 *k_B_T* to maintain an excluded volume and an *r_c_* of 2 ⋅ *r_0_*, where *r_0_* differs between the membrane and protein. Hence, *r_0_* and *B* distinguish the species in the model. The membrane model is three linear beads: the two on the ends are smaller and only interact with other lipids while the central membrane bead is larger and attracted to other membrane beads as well as the membrane binding interface of the protein. The smaller beads have a B value of 0 (i.e., no attraction), and an *r_0_* value of 1.125 nm. The central membrane beads have a membrane-membrane a *B* value of 0.6 *k_B_T* and *r_0_* value of 1.5 nm. The PS-like and PIP_2_-like membrane beads interact the same with each other, the key difference being the protein-membrane interaction. With this parameter set, the quasi-monolayer is fluid with a bending modulus around 10 *k_B_T*. The protein model is made of three curved of beads to capture the curvature and size of the I-BAR domain of IRSp53 with a *r_0_* value of 1.7 nm. There are two outer strings of beads that capture the shape with *B* value of 0 while the central string is the membrane binding interface and is attracted to the PS-like membrane beads with a *B* value of 0.235 *k_B_T* and the ends of the membrane binding interface have *B* value of 0.235 *k_B_T* to PS-like membrane beads and *B* value of 0.705 *k_B_T* to PIP_2_-like membrane beads. The PIP_2_-like membrane beads are attracted to the ends of the I-BAR domain model to recapitulate the experimentally observed behaviour of PIP_2_ clustering by I-BAR domains (*32*).

All systems were run using the LAMMPS molecular dynamics engine with a timestep of 200 fs (*77*). The initial positions of the system were a flat membrane of 188,031 membrane beads with 65 I-BAR proteins slightly above the membrane. The systems were equilibrated for 50 million timesteps in the NP_XY_L_Z_T ensemble at 0 surface tension using the Parrinello-Rahman barostat with dampening constant of 200 ps (*78*). Temperature was maintained using the Langevin thermostat with dampening constant of 20 ps (*79*). The simulations were then ran for another 500 million timesteps in the NVT ensemble.

Clustering analysis was performed by creating a network of neighbors, where I-BAR domains were neighbors if they were within 2 nm in the xy-plane. Two I-BAR domains were part of the same cluster if and only if a path existed in the network of neighbors. Thus, any I-BAR domain without a neighbour was considered free and not part of any aggregate. The probability density was estimated using kernel density estimation from the histogram of neighbour sizes from the last 250 million timesteps of the simulation and subsequently used to calculate the probability of I-BAR domain aggregates containing less than five I-BAR domains and five or more I-BAR domains. The PIP_2_-like membrane beads were separated into two groups: within 2 nm of an I-BAR domain in the xy-plane (neighboring membrane beads) and not within 2 nm of an I-BAR domain (bulk membrane beads). The percent of PIP_2_-like membrane beads within each group was quantified to measure the enrichment of PIP_2_ near I-BAR domains. The CG model analysis was performed using numpy, scikit-learn, and freud python packages and results plotted using matplotlib python package (*80–83*). All snapshots of the CG systems were created using VMD 1.9.2 (*84*).

### Live cell imaging experiments

#### Cells and transfection

Rat2 fibroblasts (ATCC, CRL-1764) were cultured in DMEM-high glucose media supplemented with 10% FBS and penicillin-streptomycin-glutamine. Cells were maintained in an incubator with 5% CO2, at +37^0^C temperature. For live cell imaging, the following constructs were used: pEGFP-N1-IRSp53L, pmTagRFP-VASP, fascin-pmCherry and empty-pmCherry. Cells were transiently transfected one day prior imaging using Xfect^TM^ transfection reagent (Takeda) according to the manufactureŕs recommendations and replated on fibronectin-coated glass-bottomed dishes (# 81158, Ibidi) two hours prior to imaging.

#### Live cell imaging

Time-lapse image series for Rat2 cells transfected with either pEGFP-N1-IRSp53L/pmTagRFP-VASP or pEGFP-N1-IRSp53L/fascin-pmCherry, or pEGFP-N1-IRSp53L/empty-pmCherry were obtained using Widefield microscope GE DeltaVision Ultra with extra sensitive cameras and Solid State Illuminators (SSI) for fluorescence excitation, equipped with incubation system set at +37^0^C and 5% CO2 and 63x or 100x oil objectives. Initial deconvolution of acquired time-lapse series were done using build-in microscopy software (softWoRx^TM^). Further analysis was conducted with Microscopy Image Browser (MIB, free MatLab-based software developed by Ilya Belevich).

#### Quantification and statistical analysis

Each analysed filopodia was monitored from the initiation of membrane bending (cone formation) until the beginning of active retraction. Changes in spatial coordinates of the positions of base and tip with the times were used to calculate the length of growing filopodia (“Distance”). The rate of growth (µm/sec) was calculated as the first derivative of filopodia length with respect to time 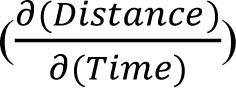. Changes in rate of growth were used to assess peculiarities of filopodia formation (stops, when 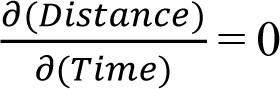 and instantaneous retractions, when 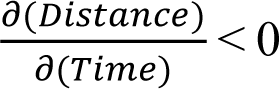). Frequency of each event (sec^-1^) was calculated as number of stops or retractions per second registered for the period of active filopodia growth.

Statistical differences in the rate of filopodia growth and the frequency of filopodia retractions were assessed by the Mann-Whitney nonparametric test at level 0.05. The statistical parameters (number of filopodia and p-values) are listed in the figure legends. Box charts are done with OriginPro 2020 program (Whiskers range within 1.5 IQR with outliers).

To assess the effect of fascin accumulation in filopodia on their elongation rates, separate kymographs of growing filopodia (time-space plot) for IRSp53-GFP and fascin-mCherry channels were generated using Fiji/ImageJ 1.53i (Multi-Kymograph plug-in, https://www.embl.de/eamnet/html/kymograph.html). Velocities “before fascin” and “with fascin” were measured for selected linear intervals of the kymographs (about 10-20 sec before (1) and after (2) fascin appearance) using ImageJ macro (tsp040221.txt). Results obtained from 26 analysed filopodia were presented as line series.

### Membrane nanotube pulling experiments

#### Cell culture

Rat2 fibroblasts (ATCC) were cultured at 37°C in DMEM GlutaMAX (Gibco, Thermo Fisher Scientific, Waltham, Massachusetts) supplemented with 1 mM sodium pyruvate (Gibco), 10% fetal bovine serum (Eurobio, Les Ulis, France), and 1% penicillin and streptomycin (Sigma-Aldrich, St. Louis, Missouri). Transient transfection of Rat2 cells was performed with FuGENE HD (Promega, Madison, Wisconsin) according to the manufacturer’s protocol. Stable cell lines were generated using G418 (geneticin) selection (0.5 mg mL^-1^) and fluorescent positive cells were sorted using a SH800 cell sorter (Sony Biotechnology, San Jose, California). Rat2 cells were routinely monitored for mycoplasma contamination and found to be negative.

#### Bead preparation

An aliquot of streptavidin-coated polystyrene beads (SVP-30-5, 0.5% w/v, Spherotech, Lake Forest, Illinois) having a nominal diameter of 3 μm was washed three times in a 10X volume of phosphate-buffered saline (PBS). Between washing steps, beads were pelleted using a centrifuge at 12000 rpm for 5 min. The final pellet was resuspended in PBS to a concentration of 0.05% w/v. Next, a volume of biotin-conjugated concanavalin A (ConA; C2272, Sigma-Aldrich, St. Louis, Missouri), having a stock concentration of 1 mg mL^-1^ in PBS, was added to the bead suspension assuming a binding capacity of 10 μg protein per mg of solid particles. The mixture was incubated overnight at 4°C on a tabletop shaker set to 1500 rpm. The ConA-coated beads were rinsed three times according to the steps above and finally resuspended in PBS to a concentration of 0.5% w/v. ConA-beads were stored at 4°C and generally usable up to one month.

#### Optical tweezer setup

A custom-built optical tweezer setup coupled to an inverted Nikon C1 Plus confocal microscope (Tokyo, Japan), as previously described in (*57*), was used for pulling plasma membrane nanotubes, force measurements, and simultaneous fluorescence imaging. Briefly, a 1064 nm continuous wave Ytterbium fiber laser (IPG Photonics, Oxford, Massachusetts) set to a 3 W input power was modulated to 400 mW (measured at the back aperture of the objective) using a polarizing beam splitter (Thorlabs, Newton, New Jersey), expanded through a telescope consisting of two plano-convex lenses with focal lengths of 100 mm and 150 mm (Thorlabs), and directed towards the back aperture of a Nikon CFI Plan Apochromat Lambda 100X 1.45 NA oil immersion objective (Tokyo, Japan). The trap stiffness *κ* was determined using the viscous drag method, including Faxen’s correction for calibration near surfaces (*85*), and averaged 60 pN μm^-1^. Displacements of a trapped bead from the fixed trap center were recorded using an Allied-Vision Marlin F-046B CCD camera (Stadtroda, Germany) at a frame rate of 20 frames per second, and later analyzed by a custom MATLAB (Mathworks) script utilizing the *imfindcircles* function to output the center location of the tracked bead (in μm). Forces were calculated from the determined bead positions according to the equation, *F* = *κ* · (*x* − *x*_o_), where *κ* is the trap stiffness, *x* is the displaced bead position, and *x*_o_ is the equilibrium reference position of the trapped bead. As the optical trap itself was stationary, all relative movements were performed using a piezo-driven stage (Nano-LP100, MadCityLabs, Madison Wisconsin). Atop the stage, a temperature and CO_2_ controllable Tokai Hit STXG-WELSX stage-top incubator (Gendoji-cho, Japan) was attached, allowing cells to be maintained at 37°C in a humidified, 5% CO_2_ atmosphere during experimentation.

Confocal images were acquired using solid-state excitation lasers: a 488 (Coherent, Santa Clara, California) and a 642 nm (Melles Grot, Carlsbad, California). The detection pathway consisted of a T560lpxr dichroic beamsplitter (Chroma, Bellows Falls, Vermont), a ET525/50 bandpass filter (Chroma), a ET665 longpass filter (Chroma), and two single-photon avalanche diodes (τ-SPAD, PicoQuant, Berlin, Germany). The τ-SPADs were controlled by the SymPhoTime 64 software (PicoQuant).

#### Nanotube pulling experiments

The day before, Rat2 cells expressing I-BAR eGFP or IRSp53 FL eGFP were plated on fibronectin-coated (35 μg mL^-1^) glass bottom dishes (35 mm, No. 1.5, MatTek, Ashland, Massachusetts) at a density of ∼30,000 cells cm^-2^. Thirty to sixty minutes prior to experimentation the phenol-containing culture medium was removed, cells were rinsed with PBS, and phenol-free DMEM containing ProLong^TM^ Live Antifade Reagent (Invitrogen, Carlsbad, California) at a 1:75 dilution, and 2 mg mL^-1^ β-Casein (>98% pure, from bovine milk, Sigma-Aldrich), for surface passivation, was applied. The cells were taken to the optical tweezer setup and labeled with Cell Mask^TM^ Deep Red plasma membrane stain (Invitrogen) at a 1:2000 dilution for 10 minutes, and ConA-coated beads were added (1:50 – 1:100 dilution). Using a custom LabVIEW (National Instruments, Austin, Texas) program to control the piezo stage, membrane nanotubes were pulled by trapping an isolated floating bead, bringing it into contact with the cell for a short period of time (<10 sec), and then moving the cell away from the bead in the x direction. Confocal images encompassing the nanotube and some of the cell body (typically 1024 x 512 pixels, 5X zoom) were gathered 5 minutes after the nanotube was pulled for protein sorting analysis; identical acquisition parameters were used when gathering the membrane and protein channel data for an individual nanotube.

### Sorting analysis

Image analysis was performed using custom-written macros in the ImageJ software. To quantify protein enrichment in the nanotube (*t*) relative to the bulk plasma membrane of the cell (*ref*), the sorting parameter (*S*) is defined as 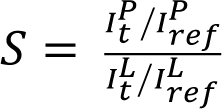 where the ratio of the green protein fluorescence (I^P^) in the nanotube and in the cell is normalized by the same ratio of the red lipid fluorescence (I^L^). This is to correct for cell-to-cell differences in both the protein and membrane signal intensity (e.g., protein expression levels, different acquisition parameters between cells, efficiency of membrane staining). Sorting values of *S* > 1 signify the protein is enriched in the nanotube while *S* < 1 signify the protein is excluded from the nanotube, with respect to its average density in the cell plasma membrane. Given Rat2 fibroblasts are quite flat, we focused on the ventral side of the cell to obtain reference images of the plasma membrane for both the protein and membrane channels. Images were background subtracted and common regions of interest (∼20,000 pixels in size) were then manually drawn to encompass homogeneous areas of the plasma membrane (near the site of the nanotube and excluding areas where vesicular puncta were observed) to calculate average protein and lipid reference values, *I^P^_ref_* and *I^L^_ref_*, respectively. The protein and membrane channels of the in-focus nanotube were also subjected to background subtraction and then normalized by their respective reference values to generate a heat map of *S* values. The resulting *S* map was filtered by a 3×3 adaptive median filter (https://weisongzhao.github.io/AdaptiveMedian.imagej/) to remove spurious pixels in the background; this processing was done given the pixel values of the raw images are discrete values (photon counts, cts) and not continuous values. The width of the nanotube was fit to a Gaussian and a rectangular region of interest (ROI; size ±2σ of the Gaussian profile) was defined along the length of the nanotube in the *S* map. Orthogonal cross sections were iteratively generated pixel-by-pixel along the length of the nanotube within the ROI. The maximum *S* value for each cross section was determined and then averaged to report the mean sorting value of the protein in the nanotube (*S_avg_*).

## Acknowledgments

We thank Evelyne Coudrier and Camille Simon for insightful discussions. We also thank Fahima Di Federico for handling plasmids, Fanny Tabarin-Cayrac for cell sorting and Anne-Sophie Mace for ImageJ programming assistance. F-CT, CLC and PB are members of the CNRS consortium AQV. F-CT and PB are members of the Labex Cell(n)Scale (ANR-11-LABX0038) and Paris Sciences et Lettres (ANR-10-IDEX-0001-02). The authors greatly acknowledge the Cell and Tissue Imaging core facility (PICT IBiSA), Institut Curie, member of the French National Research Infrastructure France-BioImaging (ANR10-INBS-04).

## Funding

Human Frontier Science Program (HFSP) grant RGP0005/2016 (F-CT, JMH, GAV, PL and PB) Institut Curie and the Centre National de la Recherche Scientifique (CNRS) (F-CT, JMH, and PB) Marie Curie actions H2020-MSCA-IF-2014 (F-CT) EMBO Long-Term fellowship ALTF 1527-2014 (F-CT) Pasteur Foundation Fellowship (JMH) Agence Nationale pour la Recherche ANR-20-CE13-0032 (JMH and PB) Université Paris Sciences et Lettres-QLife Institute ANR-17-CONV-0005 Q-LIFE (PB) FY 2015 Researcher Exchange Program between the Japan Society for the Promotion of Science and Academy of Finland (YS) Takeda Science Foundation (YS) Agence Nationale pour la Recherche ANR-18-CE13-0026-01 and ANR-21-CE13-0010-03 (CLC) Cancer Society Finland 4705949 (PL) United States National Institutes of Health (NIH) Institute of General Medical Sciences (NIGMS) grant R01-GM063796 (GAV)

## Author contributions

GAV, PL and PB designed the initial project. F-CT, JMH, ZJ, EK, YS, JP, OM, JM performed experiments and analyzed results under the supervision of CLC, GAV, PL and PB. F-CT and JP developed GUV experiments, and F-CT performed and analyzed data from the GUV experiments. JMH, ZJ, EK and JP performed and analyzed data from nanotube pulling experiments, computer simulations, live-cell imaging experiments, and actin pyrene assays, respectively. YS, JP and JM purified proteins. OM analyzed protein clustering on GUVs. F-CT wrote the original draft with inputs and revisions from JMH and PB. Conceptualization: F-CT, JMH, GAV, PL, PB Methodology: F-CT, JMH, ZJ, JP Investigation: F-CT, JMH, ZJ, EK, JP Resources: YS, JP, JM Visualisation: F-CT, JMH, ZJ, EK Supervision: CLC, GAV, PL, PB Writing—original draft: F-CT, JMH, PB Writing—review & editing: F-CT, JMH, ZJ, EK, YS, JP, OM, JM, CLC, GAV, PL, PB Funding acquisition: GAV, PL, PB

## Competing interests

The authors declare that they have no competing interests.

## Data and materials availability

All data needed to evaluate the conclusions in the paper are available in the main text and the Supplementary Materials.

## Supplementary Materials

**Fig. S1.**
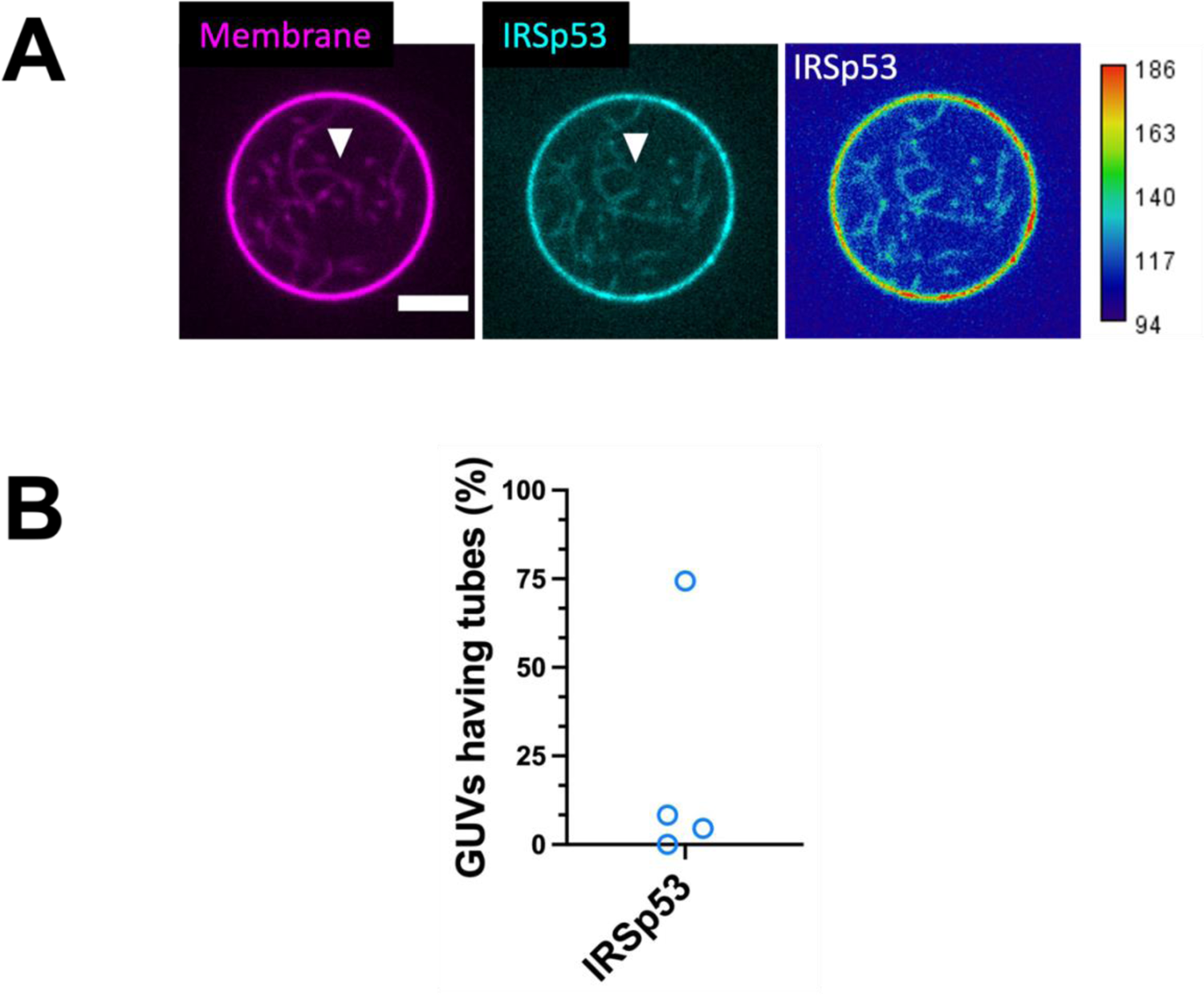
IRSp53 generates inward membrane tubes on GUVs. (A) Representative confocal images of GUVs incubated with AX488 labelled IRSp53 (16 nM, dimer). Color codes: magenta, membrane and cyan, IRSp53. Heat map: high IRSp53 intensity in red and low intensity in blue. White arrowhead indicates some membrane tubes. Scale bar, 5 μm. (B) Percentages of GUVs having tubes generated by AX488 labelled IRSp53 (16 nM, dimer). N = 22, 24, 43, 15 GUVs, n = 4 sample preparations.

**Fig. S2.**
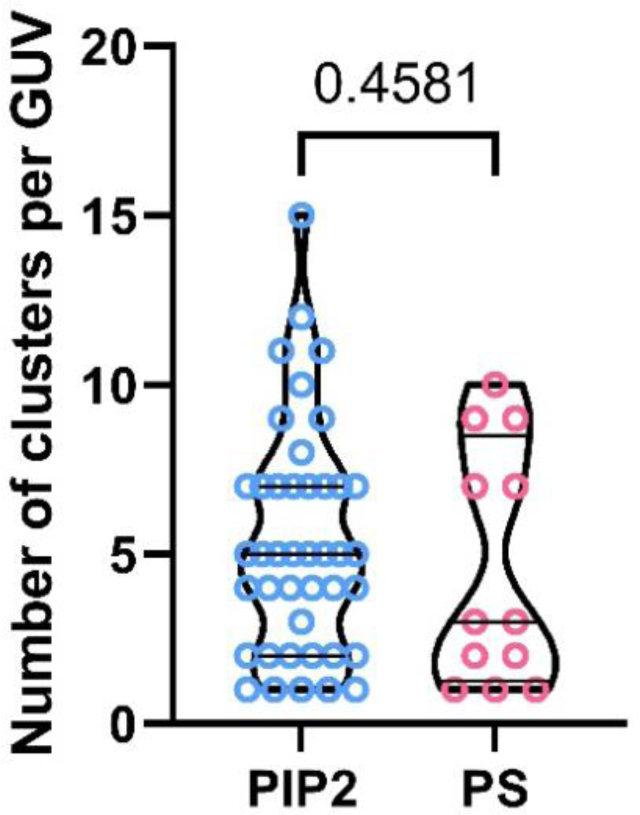
Comparable numbers of IRSp53 clusters on PIP_2_-GUVs and PS-GUVs. Characterization of the numbers of IRSp53 clusters on PIP_2_-GUVs (“PIP2”) and PS-GUVs (“PS”). “PIP2”, N = 225 clusters from 58 GUVs, n = 3 sample preparations. “PS”, N = 55 clusters from 20 GUVs, n = 2 sample preparations. Statistical test: two-tailed Mann-Whitney test, *p* = 0.4581.

**Fig. S3.**
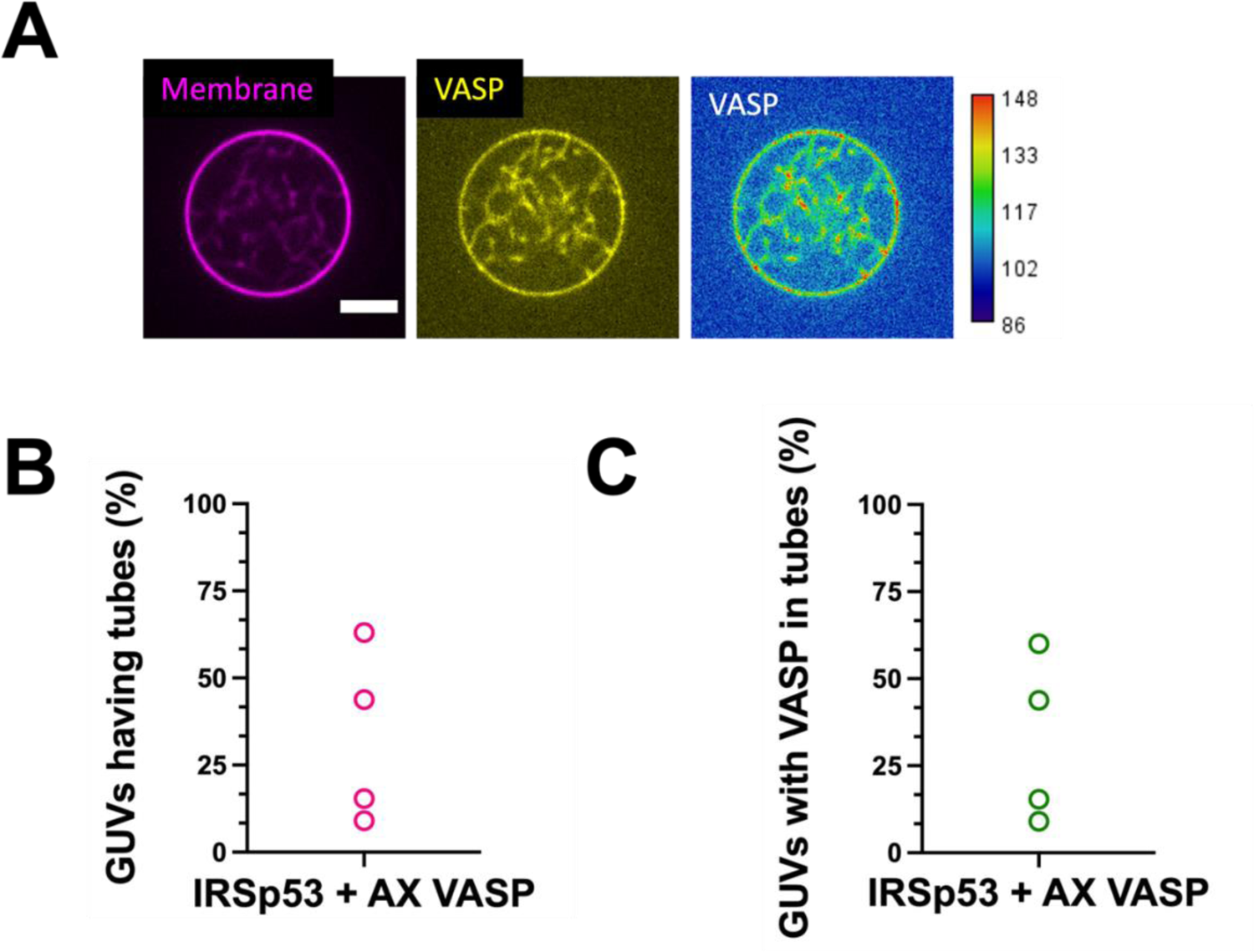
VASP is recruited to IRSp53 generated membrane tubes on GUVs. (A) Representative confocal images of GUVs incubated with IRSp53 (unlabeled, 16 nM dimer) and AX488 labelled VASP (4 nM tetramer). Color codes: magenta, membrane and yellow, VASP. Heat map: high VASP intensity in red and low intensity in blue. Scale bar, 5 μm. (B) Percentages of GUVs having tubes in the presence of unlabeled IRSp53 (16 nM, dimer) and AX488 labelled VASP (4 nM, tetramer). N = 16, 13, 43, 11 GUVs, n = 4 sample preparations. (C) Percentages of GUVs having VASP-positive tubes in the presence of unlabeled IRSp53 (16 nM, dimer) and AX488 labelled VASP (4 nM tetramer). N = 16, 13, 43, 11 GUVs, n = 4 sample preparations.

**Fig. S4.**
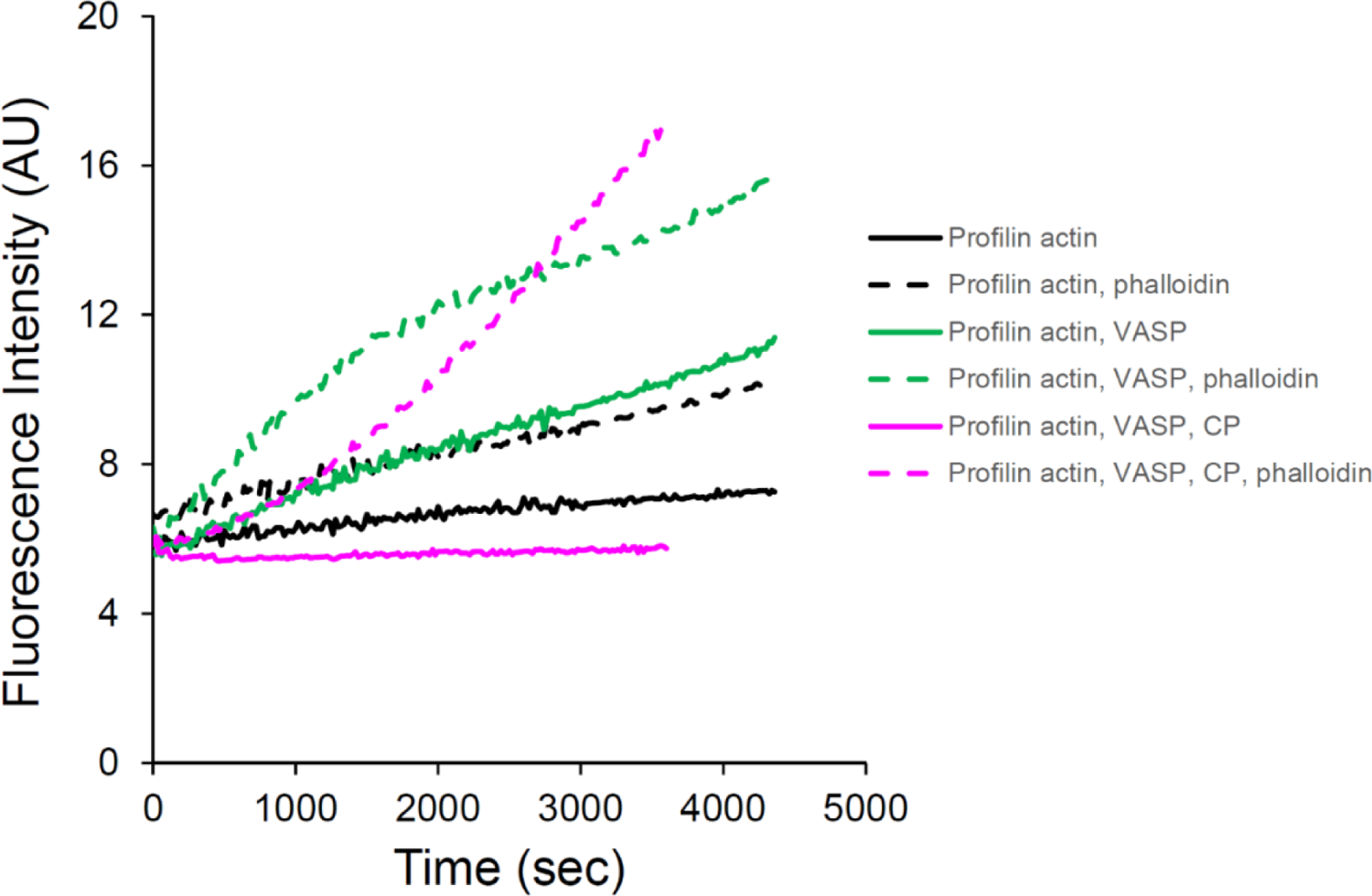
Effects of VASP, capping protein (CP) and phalloidin in profilin-actin polymerization in bulk. Pyrene actin polymerization assay. Actin (2 μM, 5% pyrenyl labeled) was spontaneously assembled with, if present, profilin (2.4 μM), VASP (15 nM in tetramer), CP (25 nM) and phalloidin (2 μM) (color coded).

**Fig. S5.**
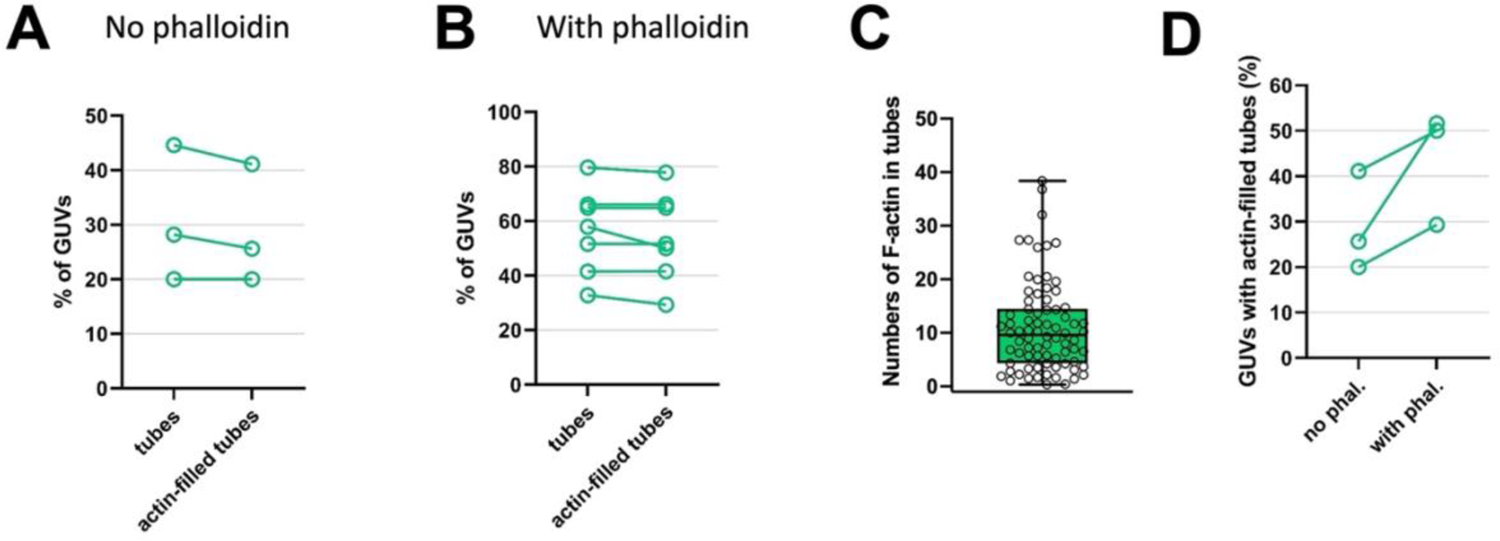
Quantifications of GUVs with actin-filled tubes and F-actin numbers in the presence and absence of phalloidin. Percentages of GUVs with membrane tubes (“tubes”) and with at least one or more actin-filled tubes (“actin-filled tubes”), (A) in the absence (“No phalloidin”) and (B) in the presence (“With phalloidin”) of AX488 phalloidin. For “No phalloidin” condition, AX488 labelled actin was used. Total GUV numbers: “No phalloidin”, 56, 39, 45; “With phalloidin”, 57, 56, 41, 54, 58, 62, 38. (C) Numbers of filamentous actin (F-actin) in tubes. N = 78 tubes, n = 3 sample preparations. (D) Total GUV numbers: In the absence of phalloidin, “no phal.”, 56, 39, 45; in the presence of phalloidin, “with phal.”, 38, 62, 58.

**Fig. S6.**
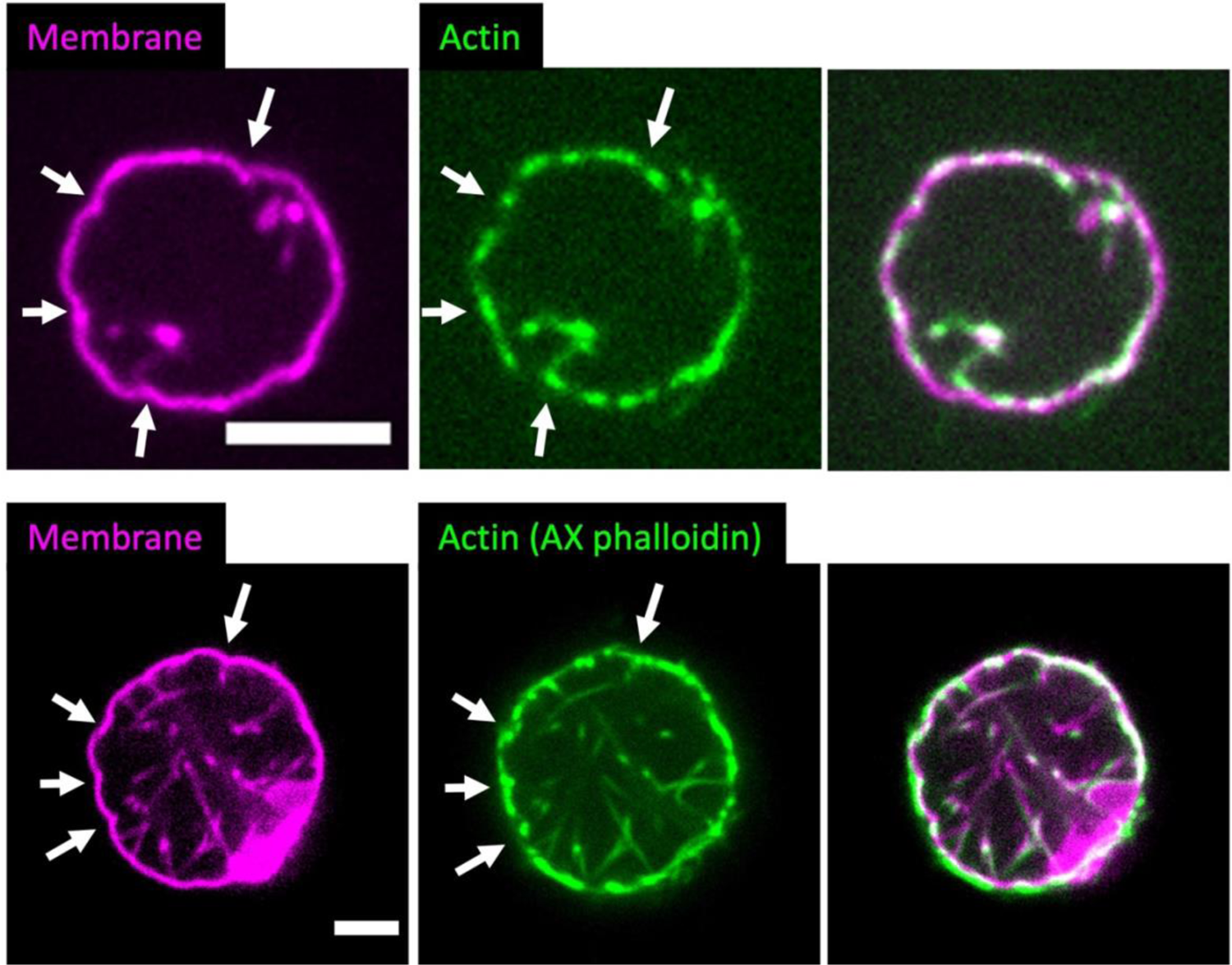
Membrane deformation by the IRSp53-VASP-actin networks. Representative confocal images of GUVs deformed by actin in the absence (Top) and presence (Bottom) of phalloidin. Magenta, membrane and green, actin. White arrows indicate some inward membrane deformations on the GUV membranes. Scale bars, 5 μm.

**Fig. S7.**
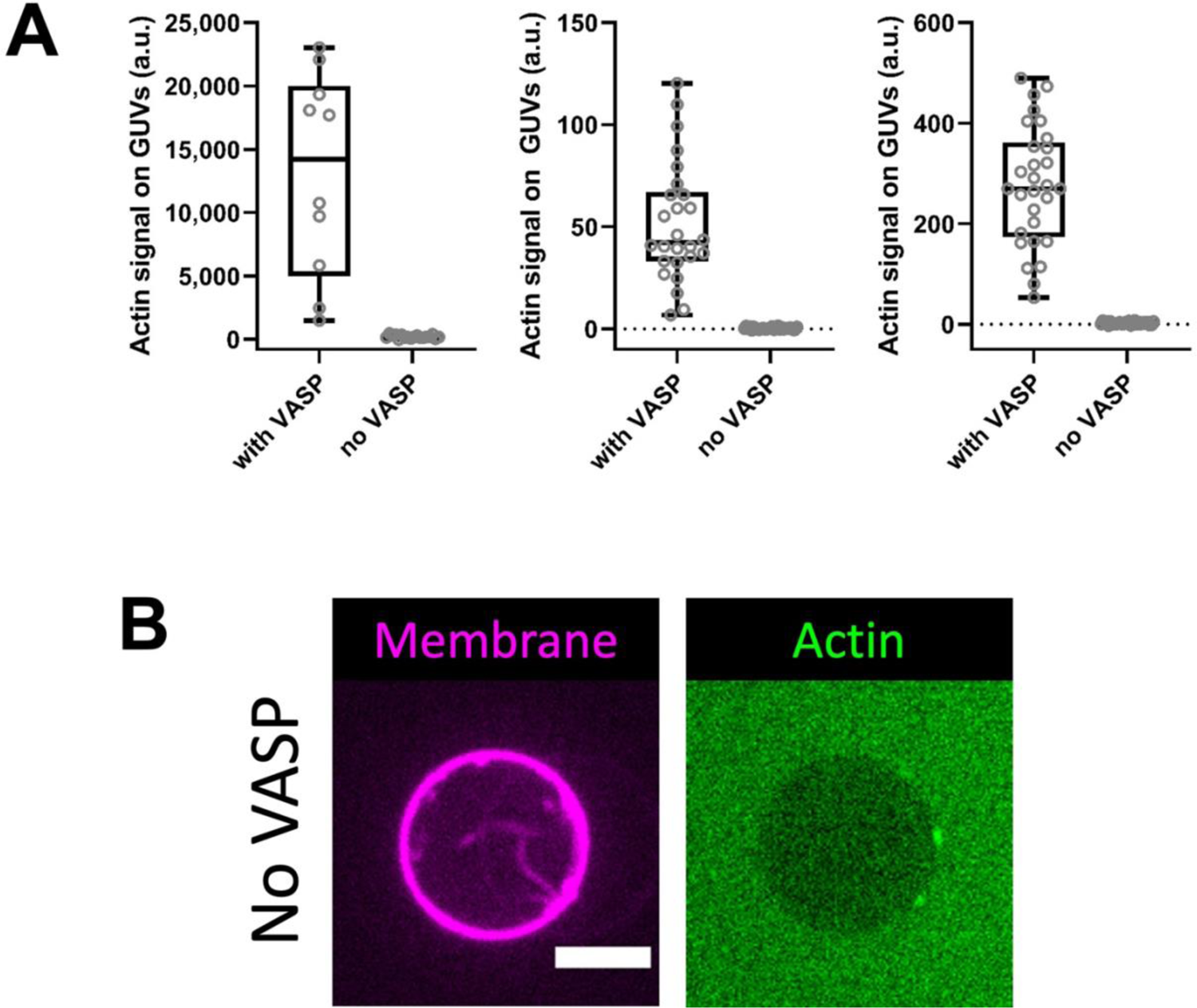
A complete lack of actin signals on GUVs in the absence of VASP. (A) The results showed three sample preparations. From left to right. N = 10, 18 GUVs, N = 26, 29 GUVs and N = 29, 29 GUVs, in the presence (“with VASP”) and absence (“No VASP”) of VASP. (B) Representative confocal images of a GUV having tubes in the absence of VASP. Membrane in magenta and actin in green. Scale bar, 5 μm.

**Fig. S8.**
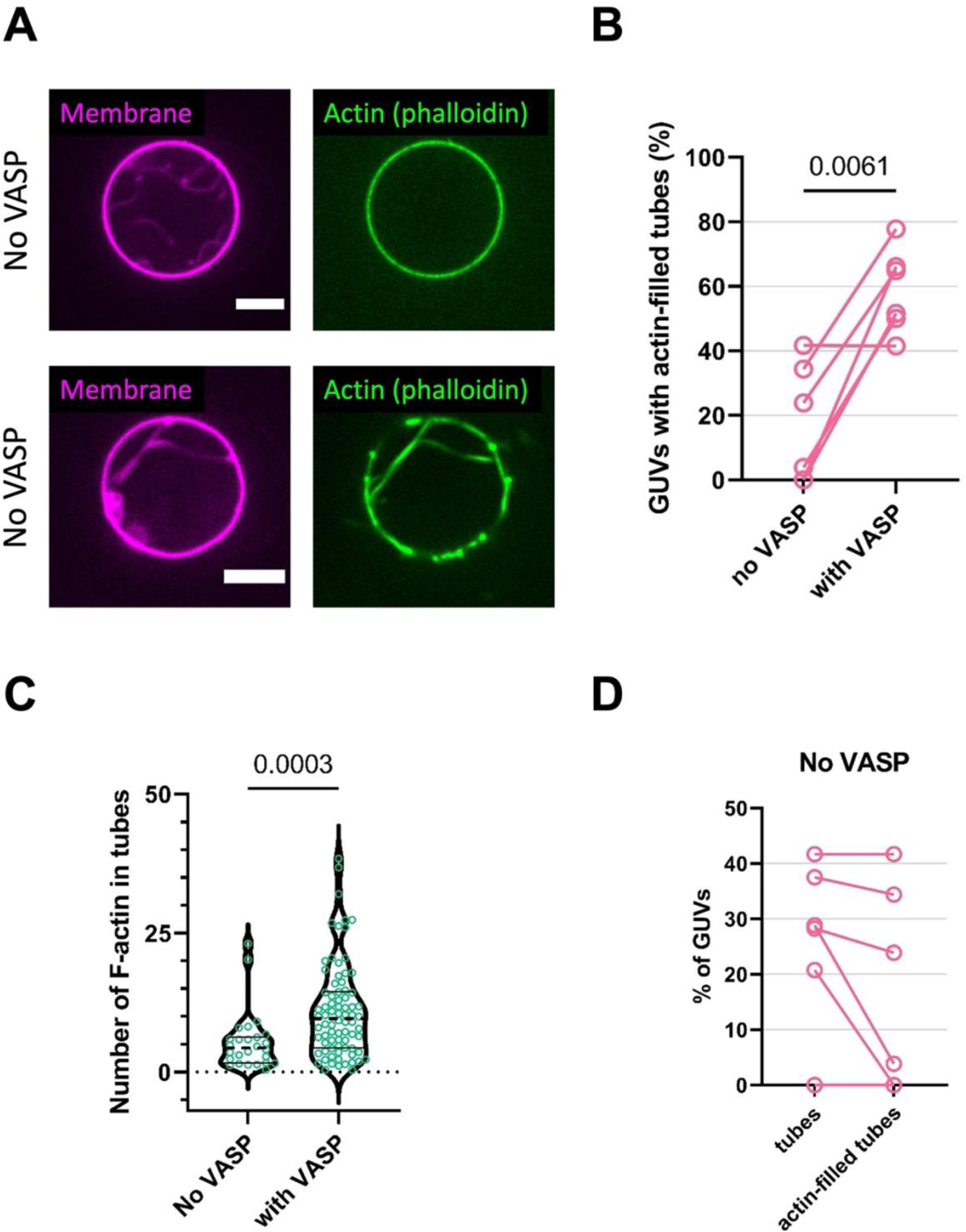
VASP facilitates protrusion formation by promoting actin polymerization in the presence of phalloidin. (A) Representative confocal images of GUVs incubated with IRSp53, actin, fascin, CP, profilin and AX488 phalloidin, i.e. in the absence of VASP. Scale bars, 5 μm. (B) Percentages of GUVs with actin-filled tubes in the absence (“No VASP”) and presence (“with VASP”) of VASP. “With VASP”, N = 41, 54, 57, 38, 56, 62 GUVs and “No VASP”, N = 24, 32, 46, 52, 11, 53 GUVs. n = 6 sample preparations. Statistical tests: (1) chi-squared test on data pooled from the 6 sample preparations, *p* < 0.0001, and (2) paired *t* test considering the 6 sample preparations individually, *p* = 0.0061 (shown in the figure). (C) Numbers of filamentous actin (F-actin) inside the tubes in the absence (“No VASP”) and presence (“with VASP”) of VASP. N = 26 (“No VASP”) and 78 (“With VASP”) tubes, n = 3 sample preparations. Note that “with VASP” data are the same as shown in Fig. S5 C. Statistical test: Mann-Whitney test, unequal variance, *p* = 0.0003. (D) In the absence of VASP, percentages of GUVs with tubes and actin-filled tubes.

**Fig. S9.**
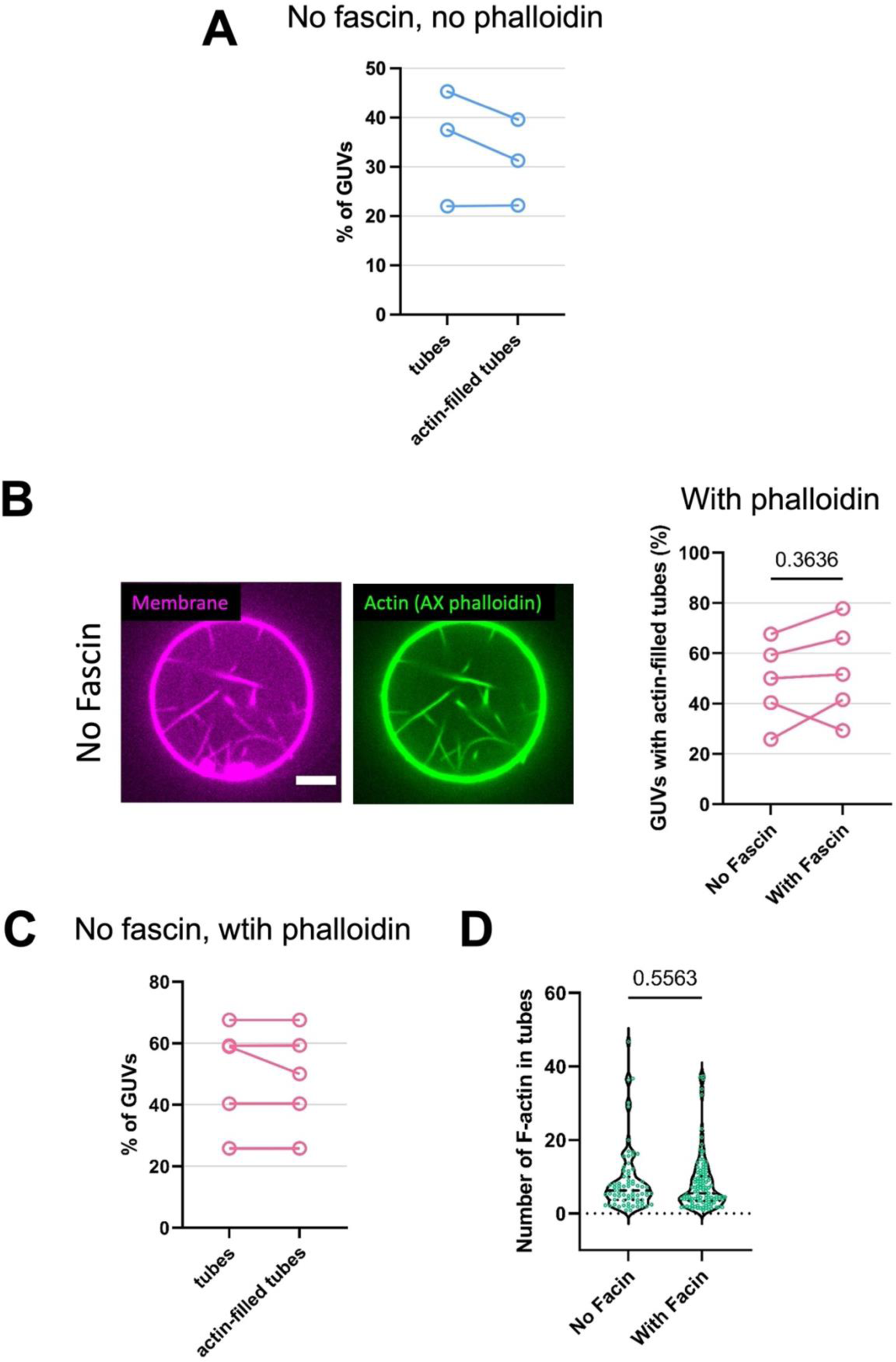
Fascin does not facilitate IRSp53-based protrusion formation. (A) In the absence of fascin and phalloidin, percentages of GUVs with membrane tubes (“tubes”) and with at least one or more actin-filled tubes (“actin-filled tubes”). To visualize actin, AX488 labelled actin was used. GUV numbers, N = 32, 54, 53, n = 3 sample preparations. (B – D) In the presence of AX488 phalloidin, (B) *Left* panel, representative confocal images of GUVs incubated with IRSp53, actin, VASP, CP, profilin and AX488 phalloidin, i.e. in the absence of fascin. Scale bars, 5 μm. *Right* panel, percentages of GUVs with actin-filled tubes in the absence (“No Fascin”) and presence (“With Fascin”) of fascin. “With Fascin”, N = 54, 56, 62, 58, 41 GUVs and “No Fascin”, N = 37, 54, 56, 57, 31 GUVs. n = 5 sample preparations. Statistical tests: (1) chi-squared test on data pooled from the 5 sample preparations, *p* = 0.3522, and (2) paired *t* test considering the 5 sample preparations individually, *p* = 0.3636 (shown in the figure). (C) In the absence of fascin, percentages of GUVs with membrane tubes (“tubes”) and with at least one or more actin-filled tubes (“actin-filled tubes”). N = 37, 54, 56, 57, 31 GUVs, n = 5 sample preparations. (D) Numbers of F-actin inside the tubes in the absence (“No Fascin”) and presence (“With Fascin”) of Fascin. N = 76 (“No Fascin”) and 109 (“With Fascin”) tubes, n = 3 sample preparations. Statistical test: Unpaired *t* test, *p* = 0.5563.

**Fig. S10.**
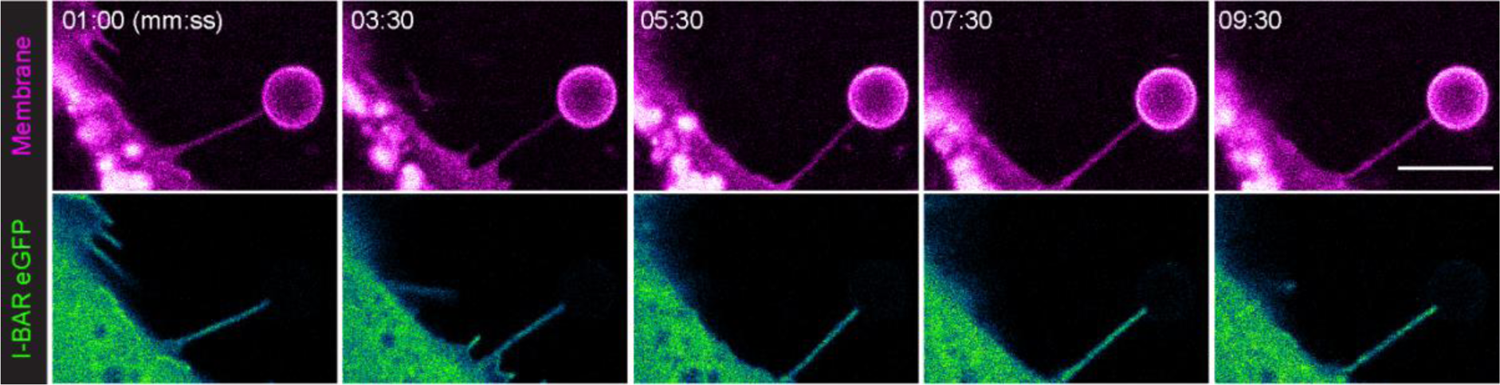
IRSp53’s I-BAR domain is visibly present in nanotubes shortly after pulling. Selected still images from Movie S8. Imaging started 1 minute after the nanotube was formed. Scale bar, 5 μm.

**Fig. S11.**
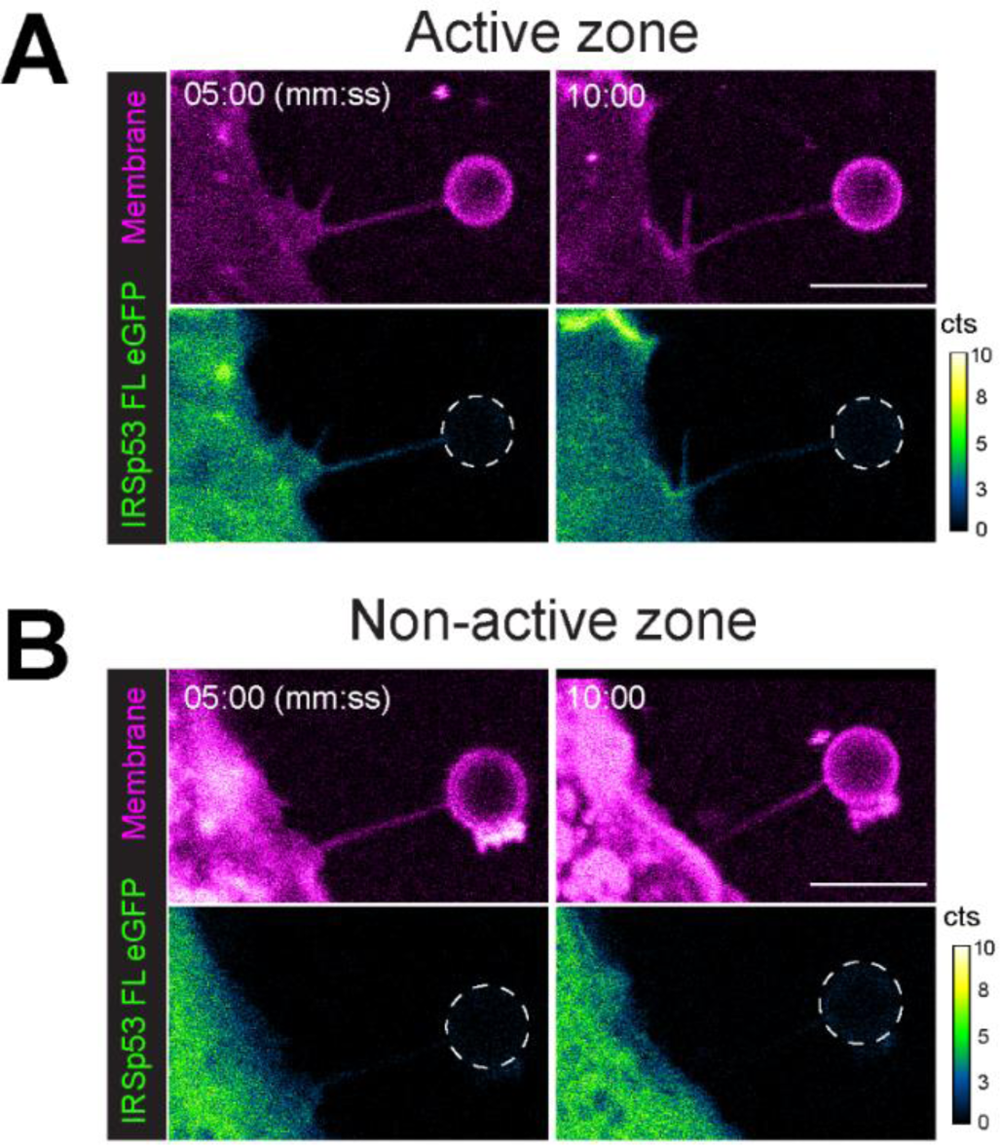
Representative nanotubes pulled from “active”. (A) and “non-active” zones (B) show that the presence or absence of IRSp53 FL, respectively, is stable through time. Indicated times are defined from the point at which the nanotube was formed. Dashed white circles outline the trapped bead. Scale bars, 5 μm.

**Fig. S12.**
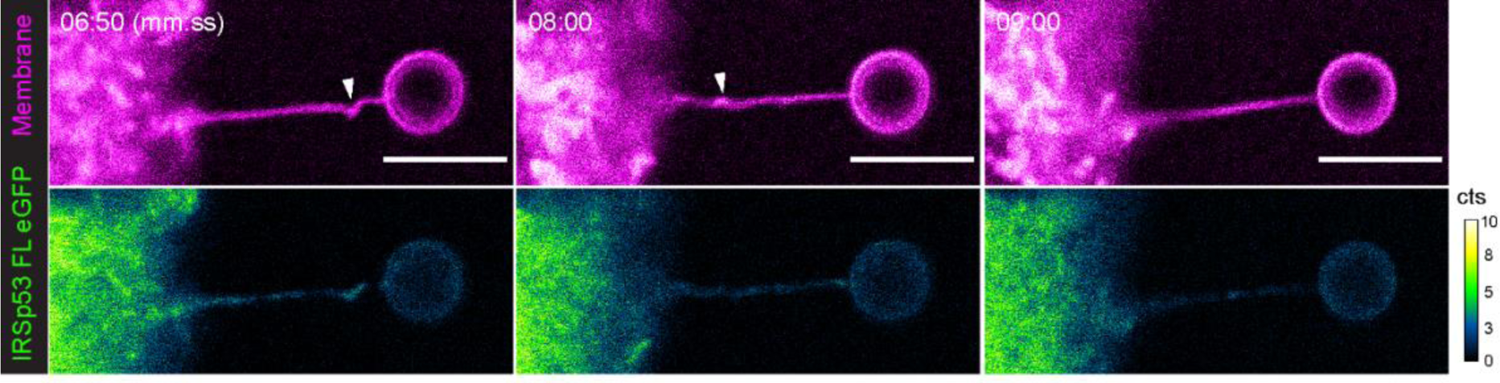
Representative example of an IRSp53 positive nanotube exhibiting helical buckling (white arrowheads). Indicated times are defined from the point at which the nanotube was formed. Scale bars, 5 μm.

---

**Movie S1. IRSp53 clusters are followed by the birth of VASP clusters at the onset of a filopodium.**

Live Rat2 cells expressing IRSp53-eGFP (green) and VASP-RFP (red). Time in seconds.

**Movie S2. IRSp53 clusters form on GUV membranes.**

Time-lapse imaging of a GUV exhibiting IRSp53 clusters. AX488-IRSp53 (cyan) at 16nM was used. GUV membranes contained 0.5 mole% of BODIPY TR ceramide (magenta). Time in mm:ss. Scale bar, 5 μm.

**Movie S3. VASP is recruited in IRSp53-generated tubes.**

Time-lapse imaging of a GUV exhibiting IRSp53-induced membrane tubes in the presence of VASP. GUV membranes contained 0.5 mole% of BODIPY TR ceramide (magenta). AX488-VASP (yellow) at 4 nM (VASP tetramer) was present together with unlabelled IRSp53 (16 nM of IRSp53 dimer). Time in mm:ss. Scale bar, 5 μm.

**Movie S4. IRSp53 drives VASP-actin-based membrane protrusions.**

Time-lapse imaging of a GUV exhibiting actin-filled membrane tubes. GUV membranes contained 0.5 mole% of BODIPY TR ceramide (magenta). A 10% - 27% of actin is tagged with AX488 (green). Time in mm:ss. Scale bar, 5 μm.

**Movie S5. IRSp53 drives VASP-actin-based membrane protrusions.**

Time-lapse imaging of a GUV exhibiting actin-filled membrane tubes. GUV membranes contained 0.5 mole% of BODIPY TR ceramide (magenta). A 10% - 27% of actin is tagged with AX488 (green). Time in mm:ss. Scale bar, 5 μm.

**Movie S6. IRSp53 drives VASP-actin-based membrane protrusions.**

Time-lapse imaging of a GUV exhibiting actin-filled membrane tubes. GUV membranes contained 0.5 mole% of BODIPY TR ceramide (magenta). AX488 phalloidin was used to visualize actin filaments (green). Time in mm:ss. Scale bar, 5 μm.

**Movie S7. IRSp53 drives VASP-actin-based membrane protrusions.**

Time-lapse imaging of a GUV exhibiting actin-filled membrane tubes. GUV membranes contained 0.5 mole% of BODIPY TR ceramide (magenta). AX488 phalloidin was used to visualize actin filaments (green). Time in mm:ss. Scale bar, 5 μm.

**Movie S8. I-BAR-eGFP is enriched in nanotubes pulled from live cells.**

Time-lapse imaging of a Rat2 cell expressing IRSp53’s I-BAR domain fused to eGFP (green). Cell Mask^TM^ Deep Red (magenta) was used to visualize the plasma membrane and the pulled nanotube. Imaging started 1 minute after the nanotube was formed. Time in mm:ss. Scale bar, 5 µm.

## Notes

### Competing Interest Statement

The authors have declared no competing interest.

